# Djp1 is a multifunctional Hsp40 cochaperone for mitochondrial phospholipid metabolism

**DOI:** 10.64898/2026.08.10.743968

**Authors:** Rashima Prem, Alex Maya-Romero, Chi Xie, Zach Irwin, Bradley Wagaman, Pingdewinde N. Sam, Salloni Gill, Ketaki Nirbhavane, Mackenzie T. Primrose, Kevin Whited, Steven M. Claypool

**Affiliations:** Department of Physiology, Pharmacology and Therapeutics, The Johns Hopkins University School of Medicine, Baltimore, MD 21205, USA; Mitochondrial Phospholipid Research Center, The Johns Hopkins University School of Medicine, Baltimore, MD 21205, USA; Department of Genetic Medicine, The Johns Hopkins University School of Medicine, Baltimore, MD 21205, USA

**Keywords:** Chaperones, protein targeting, endoplasmic reticulum, mitochondria, phospholipid metabolism

## Abstract

Mitochondria are cellular energy hubs best known for ATP production via oxidative phosphorylation; however, they also serve as biosynthetic centers for phospholipids. Mitochondrial phospholipids are critical for various cellular processes, and their loss underlies myriad mitochondrial diseases. The critical enzymes underlying these biosynthetic cascades are encoded in the nucleus, translated in the cytosol, and imported into mitochondria. Understanding of mechanisms and factors that ensure precise targeting of proteins to mitochondria has been long overlooked but remains critical. Recently, the J-protein/Hsp40 cochaperone Djp1 has emerged as a key player in mitochondrial protein targeting by promoting the transfer of precursors from the endoplasmic reticulum (ER) surface to mitochondria in a pathway termed ER-SURF. Molecular details regarding how Djp1 recognizes clients and more broadly supports mitochondrial function remain unknown. Using biochemical approaches, proteomics, and thin layer chromatography, we demonstrate that Djp1 is a regulator of Phosphatidylserine decarboxylase 1 (Psd1), an inner mitochondrial membrane resident responsible for mitochondrial phosphatidylethanolamine (PE) production. This regulation of Psd1 biogenesis is dependent on its mitochondrial targeting signal and is specific to Djp1 compared to other members of the Hsp40 family or ER targeting factors. Intriguingly, the combined loss of Djp1 and Psd1 results in a synthetic sick phenotype that unexpectedly reflects a role(s) for Djp1 in proper mitochondrial phospholipid metabolism independent of Psd1. Taken together, these findings expand our understanding of Djp1-dependent mitochondrial protein regulation and unveil Djp1 as important for mitochondrial phospholipid metabolism by multiple mechanisms.

## Introduction

Mitochondria are cellular energy hubs best known for ATP production via oxidative phosphorylation. Mitochondria also serve as biosynthetic centers for amino acids, reducing equivalents, and lipids [1]. To sustain these functions, mitochondria must import nearly 99% of their proteome which is encoded in the nucleus, translated in the cytosol and imported as precursors that are sorted into their respective compartments. Successful import requires the interaction of precursors and associated chaperones with the translocase of the outer membrane (TOM) or the mitochondrial import complex (MIM), an insertase for α-helical membrane anchored proteins on the outer mitochondrial membrane (OMM) [2–8]. At TOM, some precursors can be laterally released into the OMM [9] whereas precursors of β-barrel proteins are passed to the Sorting and Assembly Machinery prior to OMM integration [9]. Upon translocation through TOM, non-OMM destined precursors are then passed to one of two translocases of the inner membrane (TIM) before being folded and incorporated into the inner mitochondrial membrane (IMM) or instead released into the intermembrane space (IMS) or passed into the matrix [3, 7, 10].

Amongst the spectrum of nuclear encoded mitochondrial proteins are biosynthetic enzymes responsible for the synthesis of phospholipids such as phosphatidylglycerol (PG), cardiolipin (CL), and phosphatidylethanolamine (PE). These enzymes modify lipids imported from other subcellular compartments to generate phospholipids that are essential for several key mitochondrial functions including cristae formation, respiratory complex assembly and function, and mitochondrial dynamics [11–15]. The importance of proper mitochondrial phospholipid metabolism is underscored by diseases associated with mutations in key enzymes or their interaction partners [16–18]. PE is an essential membrane phospholipid produced in both the endoplasmic reticulum (ER) and mitochondrion via distinct pathways. In the ER, PE can be synthesized through lyso-PE acylation by Ale1 [19–21] or from exogenous ethanolamine via the CDP-ethanolamine Kennedy pathway [22–24]. A dedicated pool of PE in mitochondria is generated from the decarboxylation of phosphatidylserine (PS) by the IMM protein Psd1 in yeast or PISD in mammals [23, 25–27]. This nuclear encoded protein contains a bipartite N-terminus that has a mitochondrial targeting signal (MTS) followed by a hydrophobic transmembrane (TM) domain. Upon its lateral release from TIM23, membrane anchored Psd1 undergoes a self-processing autocatalytic step that results in non-covalently attached β and α subunits, the latter of which contains an N-terminal pyruvoyl group that is essential for catalytic activity [28–31]. In addition to Psd1, yeast also have Psd2, a non-conserved enzyme that resides somewhere in the endosomal system [32, 33].

Yeast lacking Psd1 exhibit impaired respiration [12, 34], growth [35], autophagic flux [36], and mitochondrial membrane fusion [13, 37]. The resultant loss of mitochondrial PE can be partially compensated for by enhanced ER-derived PE synthesis via the Kennedy pathway and in yeast, Psd2 [12]; however, limitations of these alternative sources are highlighted by the fact that loss of *PISD* is embryonically lethal in mice [38]. Moreover, *PISD* mutations in humans are associated with metabolic, neurological, and developmental defects [18, 39]. Given the demand for mitochondrial PE, it is imperative that Psd1 is efficiently targeted and imported to sustain optimal mitochondrial physiology.

Mitochondrial protein targeting refers to the process by which nuclear encoded precursors make their way to the surface of mitochondria for subsequent import. While mitochondrial protein import has been extensively studied [40], protein targeting is less understood and remains a burgeoning field. Knowledge of factors involved in protein transport to mitochondria and their associated quality control mechanisms are an overlooked aspect of mitochondrial biogenesis that has recently gained attention. The surface of the ER has emerged as a key player in mediating this critical process for a subset of mitochondrial precursors, particularly for challenging and hydrophobic substrates [7, 41–43]. The Signal Recognition Particle (SRP) is critical for the recognition and sorting of signal sequence and/or TM domain-containing precursors associated with ribosomes to the surface of the ER for co-translational import into the ER lumen [44–46]. Unexpectedly, IMM proteins Oxa1 and Psd1 were identified as SRP clients along with secretory proteins normally targeted to the ER [47]. The overlap in how proteins destined for the ER or mitochondria are targeted could increase the possibility of mistargeting. Accordingly, cells have evolved mechanisms to correct aberrant protein targeting and maintain organelle identity and function. In yeast, ER factors like Ema19 and Spf1 promote the clearance of mistargeted proteins together with the ubiquitin proteasome system [48–50]. On mitochondria, the AAA-ATPase Msp1 (ATAD1 in mammals) serves as guardian of aberrant protein targeting by recognizing mislocalized proteins and redirecting them to the ER and/or the proteasome for degradation [51–53]. Given the central role of mitochondrial function in cellular fitness, it is predictable that protein targeting involves multiple factors and layers of regulation. The importance of these mechanisms is supported by multiple lines of evidence demonstrating that improper targeting and failed import results in proteotoxicity and reduced cellular survival [53–55]. Furthermore, human ATAD1 mutations are associated with developmental and neurological defects [56–58].

Amongst the numerous recently identified mechanisms that safeguard mitochondrial protein targeting and avoid mistargeting [41, 51–54, 59, 60], the ER surface–mediated protein targeting (ER-SURF) pathway is unique in that it involves the initial targeting of mitochondrial precursors to the ER surface before engaging mitochondria. ER-SURF is mediated by Djp1, an incompletely characterized member of the J-protein/Hsp40 cochaperone family. Associated with the ER, Djp1 recognizes client precursors on the ER surface, and through ER-mitochondrial contact sites, delivers them to the TOM complex for their subsequent import [41, 42]. With only a handful of known clients [41, 42, 61], a detailed mechanistic understanding of how Djp1 recognizes its substrates and whether there are additional factors involved remain unknown. As such, it is also unresolved whether the Djp1-dependent pathway is a general targeting mechanism or a backup for “error-prone” mitochondrial precursors. Lastly, how Djp1 might coordinate with other targeting and/or quality control mechanisms to support broader mitochondrial functions like energy production or lipid metabolism also remain unknown.

Here, we demonstrate that Djp1 is critical for robust Psd1 accumulation and this is mediated through the MTS of Psd1. This Djp1 dependence is specific as evidenced by the inability of other related J-proteins to impact Psd1 steady state levels. Furthermore, we show that this regulation is necessary for proper cellular growth and cannot be fully compensated for by ER-derived PE further supporting the idea of functional divergence between ER and mitochondrial PE. Lastly, we provide evidence that Djp1 is necessary for proper mitochondrial phospholipid metabolism via a mechanism(s) unrelated to its role in promoting mitochondrial Psd1 accumulation. Taken together, this study expands our mechanistic understanding of Djp1-mediated protein targeting and uncovers Djp1 as a multifunctional protein supporting mitochondrial phospholipid metabolism.

## Results

### Djp1 post-transcriptionally controls mitochondrial Psd1 amounts regardless of metabolic state

In addition to its main localization to the inner mitochondrial membrane, a small proportion of Psd1 has been suggested to also, based on its N-glycosylation status, localize to the ER [62]. While the functional status of what was suggested to be glycosylated Psd1 was not directly tested in that study [62], it is presumably non-functional given that ER-enriched microsomal preparations lack significant Psd1 activity when the endosomal resident Psd2 is absent [33, 63]. Indeed, we recently demonstrated that in certain cellular contexts, a fraction of mutant Psd1 incapable of undergoing autocatalysis and thus devoid of PS decarboxylase activity is targeted to the ER prior to being ultimately degraded by the ubiquitin-proteasome system [64]. In the context of this Psd1-ER controversy, the Djp1-dependent ER-SURF pathway piqued our interest as a possible mechanism by which ER-associated Psd1 precursors could be directed to mitochondria for productive import. Djp1 is an ER-associated Hsp40 cochaperone modeled to associate with troublesome mitochondrial precursors before passing them to TOM for productive mitochondrial import [41, 42]. In yeast, Psd1 overexpression is sufficient to induce a cellular stress response [53], suggesting that it is a troublesome substrate to import, and the Psd1 precursor is known to engage the TOM receptors, Tom22 and Tom70 [29]. Further, the relative abundance of glycosylated Psd1 [62] and importance of the Djp1-dependent ER-SURF [41] are both suggested to be increased when the metabolic demand for mitochondrial energy production is low.

A comprehensive list of Djp1-dependent mitochondrial clients has yet to be established. As such, we hypothesized that Psd1 may be a novel ER-SURF substrate. If correct, one would expect that the total steady state amount of Psd1 would be decreased in the absence of Djp1. Consistent with our postulate, Psd1 amounts were reduced by 40-55% in yeast devoid of Djp1 regardless of yeast strain background (GA74-1A, BY4742 and W303-1A) or metabolic state ‒ glycolytic when grown in rich dextrose medium (Fig 1A) or respiratory when grown in rich lactate (Fig 1B). The absence of Djp1 did not impact the levels of *PSD1* mRNA across all three yeast backgrounds (Fig 1C), suggesting that Djp1 regulates Psd1 accumulation via a post-transcriptional mechanism. Importantly, re-introduction of Djp1 into the *djp1*Δ strain on low or high copy plasmids restored the amount of Psd1 to, but not beyond, physiologic levels (Fig 1D), indicating that the amount of Djp1 is not a limiting factor for Psd1 accumulation. To determine if the reduced amount of Psd1 detected in Djp1-lacking cell extracts reflects a decrease in its accumulation in mitochondria, subcellular fractions were collected from wildtype (WT), *djp1*Δ, and *psd1*Δ yeast grown in either rich dextrose (Fig 1E) or rich lactate (Fig 1F). Regardless of metabolic state, the amount of Psd1 that co-fractionated with the mitochondrial (P13) fraction was noticeably reduced in the absence of Djp1 (Fig 1E and 1F). As expected [41], Djp1 was enriched in the ER-containing P40 fraction. Of note, while Psd1 amounts were dramatically reduced in Djp1-lacking yeast, it still localized to mitochondria and did not accumulate in other subcellular fractions as either a mature functional protein or inactive precursor. This suggests that Djp1 does not have a role in targeting and/or inserting Psd1 into membranes of non-mitochondrial organelles. These results further highlight that the role of Djp1 in promoting the mitochondrial accumulation of Psd1 is agnostic to metabolic status, remaining high whether cellular energy is provided by glycolysis (dextrose) or oxidative phosphorylation (lactate).

**Fig 1:**
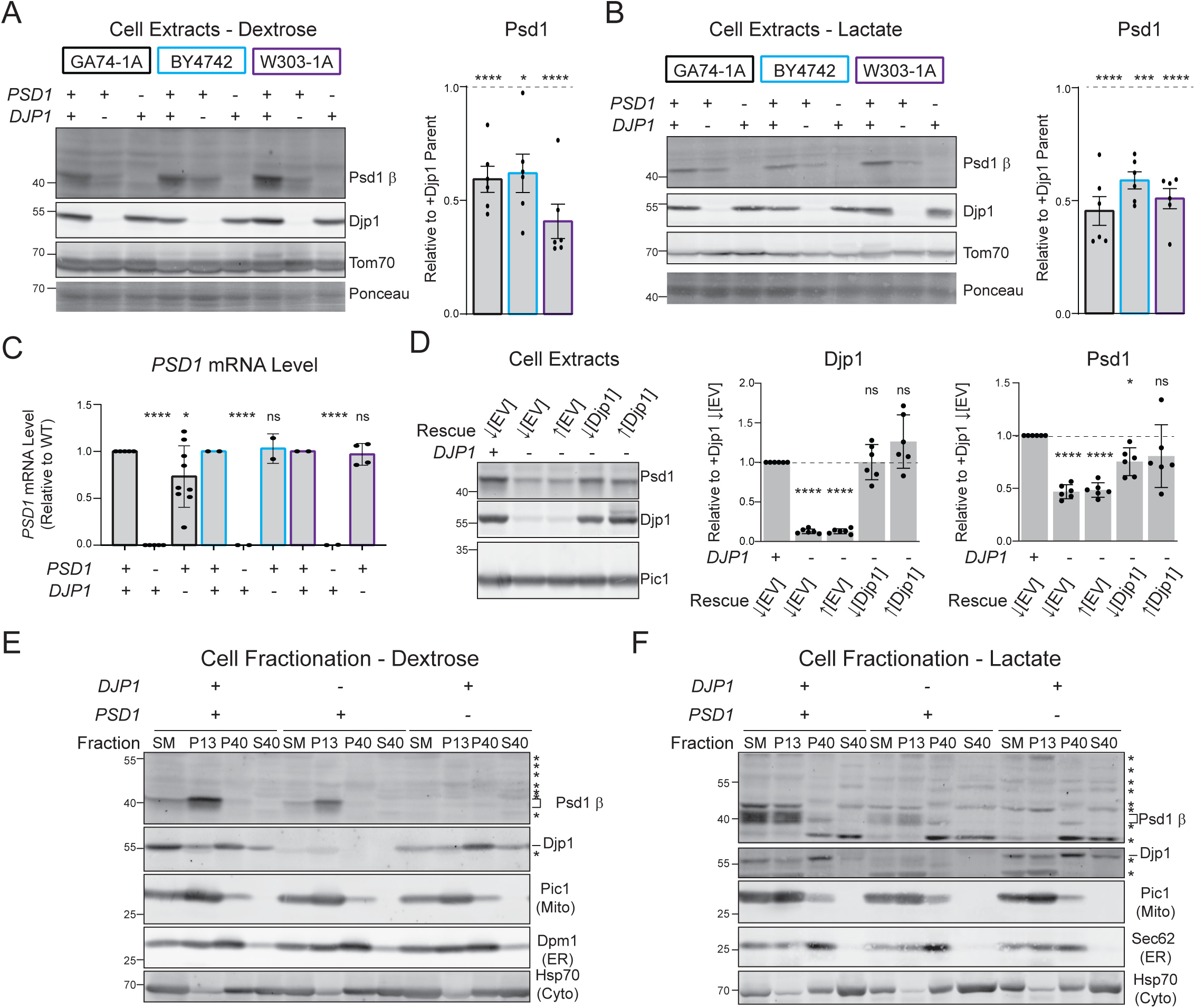
Djp1 promotes the accumulation of Psd1 in mitochondria via a post-transcriptional mechanism. **(A, B)** The Psd1 β subunit and Djp1 were detected by immunoblot in yeast cell extracts of the indicated genotype from three yeast strain backgrounds grown in rich dextrose (A) or rich lactate (B) media. Tom70 and total protein stain acted as loading controls. The relative Tom70-normalized amount of Psd1 in the absence of Djp1 versus wildtype was determined for each strain (mean ± SD for n = 6 biological replicates). **(C)** The *ACT1*-normalized *PSD1* mRNA levels in the indicated strains grown in rich lactate were determined by two-step reverse transcription quantitative PCR (mean ± SD for: GA74-1A, n = 5 WT and *psd1*Δ biological replicates and *n =* 10 biological replicates from 2 *djp1*Δ clones; BY4742, n = 2 biological replicates; and W303-1A, n = 2 WT and *psd1*Δ biological replicates and *n =* 4 biological replicates from 2 *djp1*Δ clones). For A-C, significant differences (ns, not significant; 1 symbol, *P* ≤ 0.05; 3 symbols, *P* ≤ 0.001; 4 symbols, *P* ≤ 0.0001) versus parental wildtype strains were determined by one-way ANOVA with Dunnett’s multiple comparisons. **(D)** Yeast cell extracts from WT or *djp1*Δ yeast transformed with empty vector (EV), low(↓) or high(↑)-copy Djp1 plasmids were immunoblotted as indicated. The relative Pic1-normalized amount of Djp1 and Psd1 versus wildtype [EV] was determined (mean ± SD for n = 6 biological replicates). Significant differences (ns, not significant; 1 symbol, *P* ≤ 0.05; 4 symbols, *P* ≤ 0.0001) versus wildtype [EV] were determined by one-way ANOVA with Tukey’s multiple comparisons. **(E, F)** Fractions of rich dextrose (E) or lactate (F) grown yeast of indicated genotype were harvested by differential centrifugation and equal protein amounts resolved by SDS-PAGE and immunoblotted for the Psd1 β subunit and mitochondrial (Pic1), ER (Dpm1), and cytosolic (Hsp70) controls. SM, starting material; P13, pellet of 13,000 x *g*; P40, pellet of 40,000 x *g*; S40, supernatant of 40,000 x *g*.

### Djp1 specifically controls mitochondrial Psd1 amounts

There are at least 22 Hsp40 co-chaperones in the *Saccharomyces cerevisiae* genome which are broadly categorized into three types depending upon their domain structures [65]. Type I Hsp40s have an N-terminal J-domain connected to the C-terminus client binding fragment via a phenylalanine/glycine (F/G) rich linker region and two zinc finger motifs; type II Hsp40s such as Djp1 lack the zinc finger motif in their polypeptide sequence; and type III Hsp40s do not possess any of the conserved domains apart from the essential J-domain (Fig 2A). Recently, other Hsp40 family members, including Xdj1, Ydj1, and Sis1 [66, 67], have been implicated in the handling of hydrophobic domain-containing mitochondrial precursors, suggesting a potential broader role for this family in mitochondrial biogenesis. Moreover, several yeast Hsp40 family members have been suggested to functionally overlap, a capacity that could reflect their shared structural features and/or subcellular localizations. For example, Caj1 and Djp1, both type II Hsp40s, have partially redundant roles in supporting peroxisomal protein import [68] and expression of fragments containing the J domains from multiple Hsp40 members, including Djp1, can rescue the severe *ydj1*Δ growth phenotype [69].

**Fig 2:**
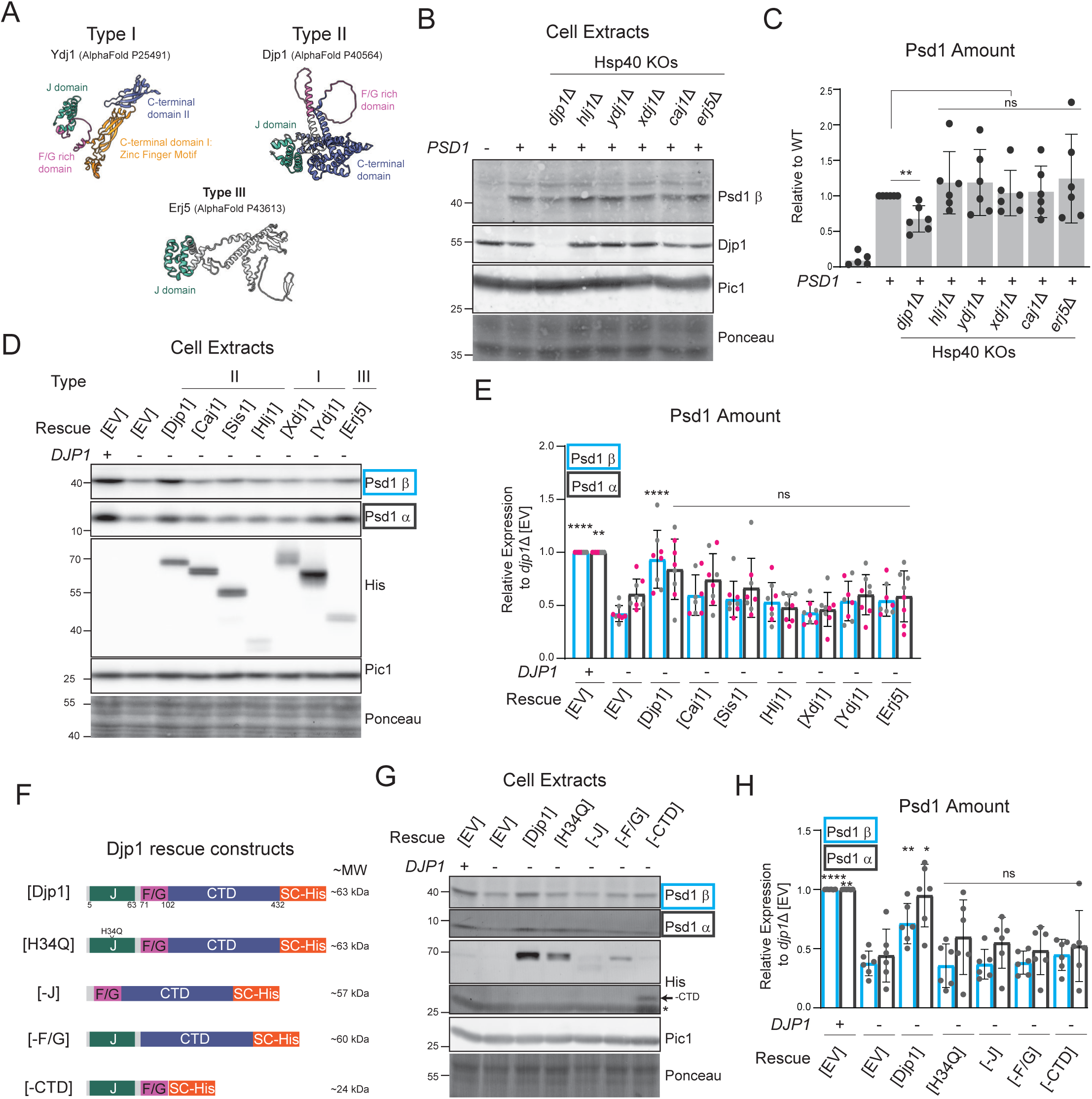
The impact on Psd1 amounts is specific to Djp1 and requires all domains. **(A)** Structural representation of the distinct domains in Hsp40 family proteins. AlphaFold images of representative Type I (Ydj1), Type II (Djp1), and Type III (Erj5) Hsp40s are shown, highlighting shared and unique domains as indicated. **(B)** The Psd1 β subunit was detected by immunoblot in yeast cell extracts of the indicated genotype grown in rich dextrose. Pic1 and total protein stain acted as loading controls. **(C)** The relative Pic1-normalized amount of Psd1 versus wildtype was determined (mean ± SD for n = 6 biological replicates). Significant differences (ns, not significant; 2 symbols, *P* ≤ 0.01) versus wildtype were determined by unpaired t tests. **(D)** Yeast cell extracts from Djp1 containing or lacking IM-Psd1 knock-in yeast (Psd1 with C-terminal 3XFLAG tag to track Psd1 α subunit) transformed with empty vector (EV) or the designated Hsp40 cochaperone were immunoblotted as indicated. Pic1 and total protein stain acted as loading controls. **(E)** The relative Pic1-normalized amount of the Psd1 β and α subunits were determined relative to wildtype [EV] (mean ± SD for n = 8 biological replicates; different clones shown in pink or gray). Significant differences (ns, not significant; 2 symbols, *P* ≤ 0.01; 4 symbols, *P* ≤ 0.0001) versus *djp1*Δ [EV] were determined by one-way ANOVA with Tukey’s multiple comparisons. **(F)** Schematic of full length, mutant, and truncated Djp1 constructs, all harboring C-terminal SpyCatcher002-8X His tags, and their predicted molecular weights (MW). **(G)** Yeast cell extracts from Djp1 containing or lacking IM-Psd1 knock-in yeast transformed with EV or the indicated Djp1 variant were immunoblotted as designated. Pic1 and total protein stain acted as loading controls. **(H)** The Pic1-normalized amount of the Psd1 β and α subunits were determined relative to WT [EV] (mean ± SD for n = 6 biological replicates). Significant differences (ns, not significant; 1 symbol, *P* ≤ 0.05; 2 symbols, *P* ≤ 0.01) versus *djp1*Δ [EV] were determined by one-way ANOVA with Tukey’s multiple comparisons.

To test whether Psd1 is an exclusive client of Djp1 or instead a shared substrate with other Hsp40 family members, we first pursued a loss-of-function strategy focusing on yeast strains lacking Hsp40 family members with documented redundancy with Djp1 (Caj1 and Ydj1), implicated in mitochondrial biogenesis (Xdj1, Ydj1, and Sis1), or that also associate with the ER (Erj5 [70] and Hlj1 [71]). The steady state abundance of Psd1 was only reduced in the *djp1*Δ strain (Fig 2B and 2C), indicating that the Psd1-Djp1 relationship is likely specific. Given that partial redundancy can sometimes only be detected upon the combined loss of genes or the overexpression of one or both participants, and that Sis1 is an essential gene, we next pursued an overexpression-based gain-of-function strategy. To this end, the ability of overexpressed Djp1, Caj1, Sis1, Hlj1, Xdj1, Ydj1, and Erj5, each containing a C-terminal SpyCatcher-His tag and under the control of the *AAC2* promoter, to restore Psd1 abundance in *djp1*Δ yeast was tested. As only Djp1 expression rescued the low-Psd1 phenotype (Fig 2D and 2E), the results from this select survey of Hsp40 members suggest that the normal accumulation of Psd1 is specifically controlled by Djp1.

Typical of type II Hsp40s, Djp1 is a 432 aa long protein that has an N-terminal J-domain required for stimulating Hsp70 ATPase activity, a middle F/G rich domain, and a C-terminal substrate binding domain. As these domains confer functional diversity and specificity upon cochaperones, we assessed their individual importance for the Psd1-Djp1 relationship by designing Djp1 constructs individually lacking each of the domains, all containing a C-terminal SpyCatcher (SC)-His tag (Fig 2F). In addition, we generated a Djp1 J domain mutant, H34Q, that is unable to stimulate Hsp70 ATPase activity. Unlike full-length Djp1, the domain-deleted and H34Q mutant Djp1 constructs all failed to rescue the low amount of Psd1 in *djp1*Δ yeast (Fig 2G and 2H). These results indicate that catalytically active full length Djp1 is required to promote the standard accumulation of Psd1.

### Psd1 amounts are unchanged or increased in absence of upstream ER-targeting pathways

We next examined whether perturbing candidate ER-targeting pathways, which could function upstream of the ER-associated Djp1, alters steady state Psd1 levels. Protein targeting to the ER can occur either co- or post-translationally. Co-translational targeting depends on SRP. Psd1 was previously identified as an SRP client by selective ribosome profiling [47]. To determine whether SRP function influences steady state Psd1 abundance, we analyzed *sec65-1*, which has a temperature-sensitive mutation in the Sec65 subunit of SRP [72]. The *sec65-1* mutant has been reported to exhibit a lethal growth phenotype at 35°C [73]. Therefore, WT and *sec65-1* cells grown at 23°C were either maintained at 23°C or shifted to the restrictive temperature of 35°C for 5 h. Following the 5-h shift to 35°C, Psd1 abundance decreased in both strains; however, no significant difference was observed between WT and *sec65-1* at either temperature (Fig 3A and 3B). This finding suggests that the SRP-dependent pathway is not required for the steady state amount of Psd1. We next turned to the guided entry of tail-anchored proteins (GET) pathway, a distinct route that mediates the post-translational ER targeting of tail-anchored proteins containing a single C-terminal transmembrane domain. GET-mediated ER targeting was recently shown to directly bind precursors of polytopic mitochondrial carriers and deliver them to the ER prior to their Djp1-assisted productive transfer to mitochondria [43]. Get3 is the cytosolic targeting ATPase that delivers GET substrates to the Get1/Get2 insertase in the ER [74]. Psd1 abundance was not significantly altered in *get3*Δ compared to WT (Fig 3C and 3D), indicating that Get3 is not required to maintain steady state Psd1 amounts.

**Fig 3.**
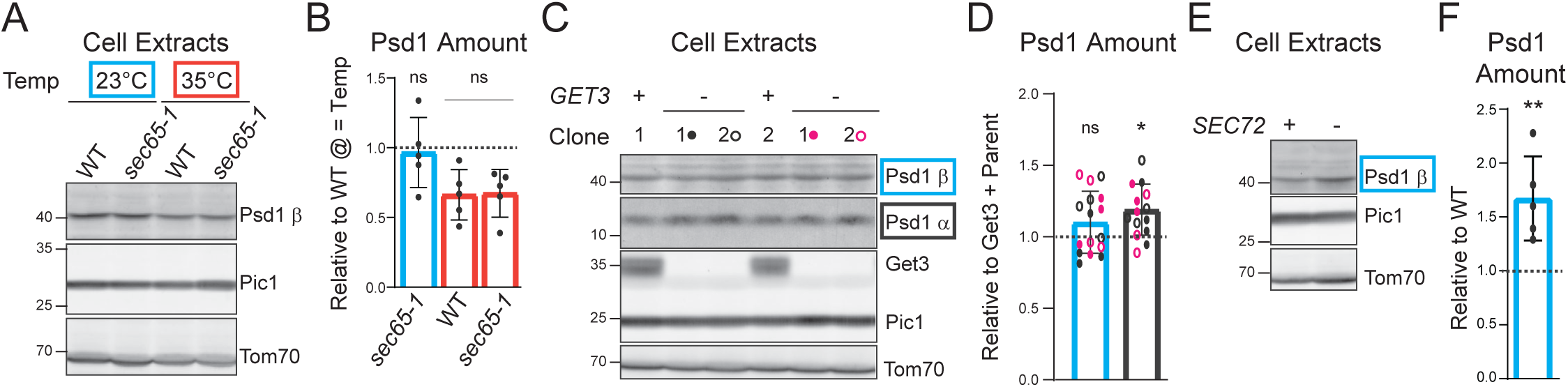
Psd1 amounts are insensitive to loss of GET or SRP-dependent pathways but increased in absence of SRP-independent pathway. **(A)** The Psd1 β subunit was detected by immunoblot in yeast cell extracts of the indicated genotype grown in rich dextrose for five hours at designated temperature. Pic1 and Tom70 acted as loading controls. **(B)** The amount of Psd1 β subunit was determined relative to WT grown at 23°C (mean ± SD for n = 5 biological replicates). Significant differences (ns, not significant) versus WT grown at corresponding temperature were tested by unpaired t tests. **(C)** Yeast cell extracts from Get3 containing or lacking IM-Psd1 knock-in yeast grown in rich lactate were immunoblotted for the Psd1 β and α subunits and Get3. Pic1 and Tom70 acted as loading controls. **(D)** The amount of Psd1 β and α subunit was determined relative to their respective Get3-containing parent (mean ± SD for n = 8 biological replicates for each Get3-positive parent/Get3-negative daughter). Significant differences (ns, not significant; 1 symbol, *P* ≤ 0.05) versus WT were tested by unpaired t tests. **(E)** The Psd1 β subunit was detected by immunoblot in yeast cell extracts of the indicated genotype grown in rich lactate. Pic1 and Tom70 acted as loading controls. **(F)** The amount of Psd1 β subunit was determined relative to WT (mean ± SD for n = 5 biological replicates). Significant differences (2 symbols, *P* ≤ 0.01) versus WT were determined by unpaired t tests.

Finally, we examined whether Psd1 abundance is influenced by Sec72-dependent SRP-independent translocation. Sec72 is a nonessential subunit of the Sec62/Sec63 complex that facilitates the translocation of a subset of SRP-independent substrates [75–77]. Loss of Sec72 diminishes but does not abolish SRP-independent translocation [78, 79]. Unexpectedly, Psd1 abundance was significantly increased in *sec72*Δ compared to WT (Fig 3E and 3F). The molecular basis for this unanticipated increase in Psd1 amounts upon loss of Sec72, and whether it reflects a direct or indirect consequence of an impaired SRP-independent pathway, is presently unclear. Collectively, these results indicate that ER-associated targeting and translocation machineries predicted to function upstream of Djp1 either do not influence (SRP-dependent and GET pathways) or surprisingly, elevate (SRP-independent pathway) the steady state Psd1 abundance.

### The MTS of Psd1 is necessary and sufficient for Djp1-dependence

With the intent to determine if the Djp1-sensitivity is intrinsic to Psd1, or perhaps reflects a general feature of mitochondrial proteins with shared features, we asked whether the abundance of IMM residents with the same IMM topology as Psd1 are also negatively impacted in absence of Djp1. Despite also being anchored in the IMM by a single TM domain with their N- and C-termini exposed to the matrix and IMS, respectively, the steady state amounts of Yme1, Mic60 and Tim50 were unaltered in Djp1-lacking cell extracts (Fig 4A). Given the apparent restricted nature of the Psd1-Djp1 relationship, we next probed where in Psd1 this specificity originates. We had previously generated a series of chimeric constructs where the N-terminal MTS and TM of Psd1 were replaced with equivalent regions of the above-mentioned IMM proteins (Fig 4B); importantly, each chimeric construct retained its self-processing activity, as evidenced by the appearance of separate Psd1 β and 3X Flag tagged Psd1 α subunits [64]. As the expression levels of Yme1, Mic60 or Tim50 are not dictated by Djp1 (Fig 4A), these constructs could help resolve whether the motif that confers Djp1-sensitivity to Psd1 is contained within the first 100 N-terminal residues, the 400 amino acids that follow the TM domain, or perhaps both. In contrast to intact Psd1 where both subunits were reduced in absence of Djp1, the steady state amounts of both subunits (the chimeric β and released α subunits) for each chimera were unchanged in the absence versus the presence of Djp1 (Fig 4 C and 4D). This indicates that the Djp1-sensitivity motif resides in the first 100 amino acids of Psd1.

**Fig 4.**
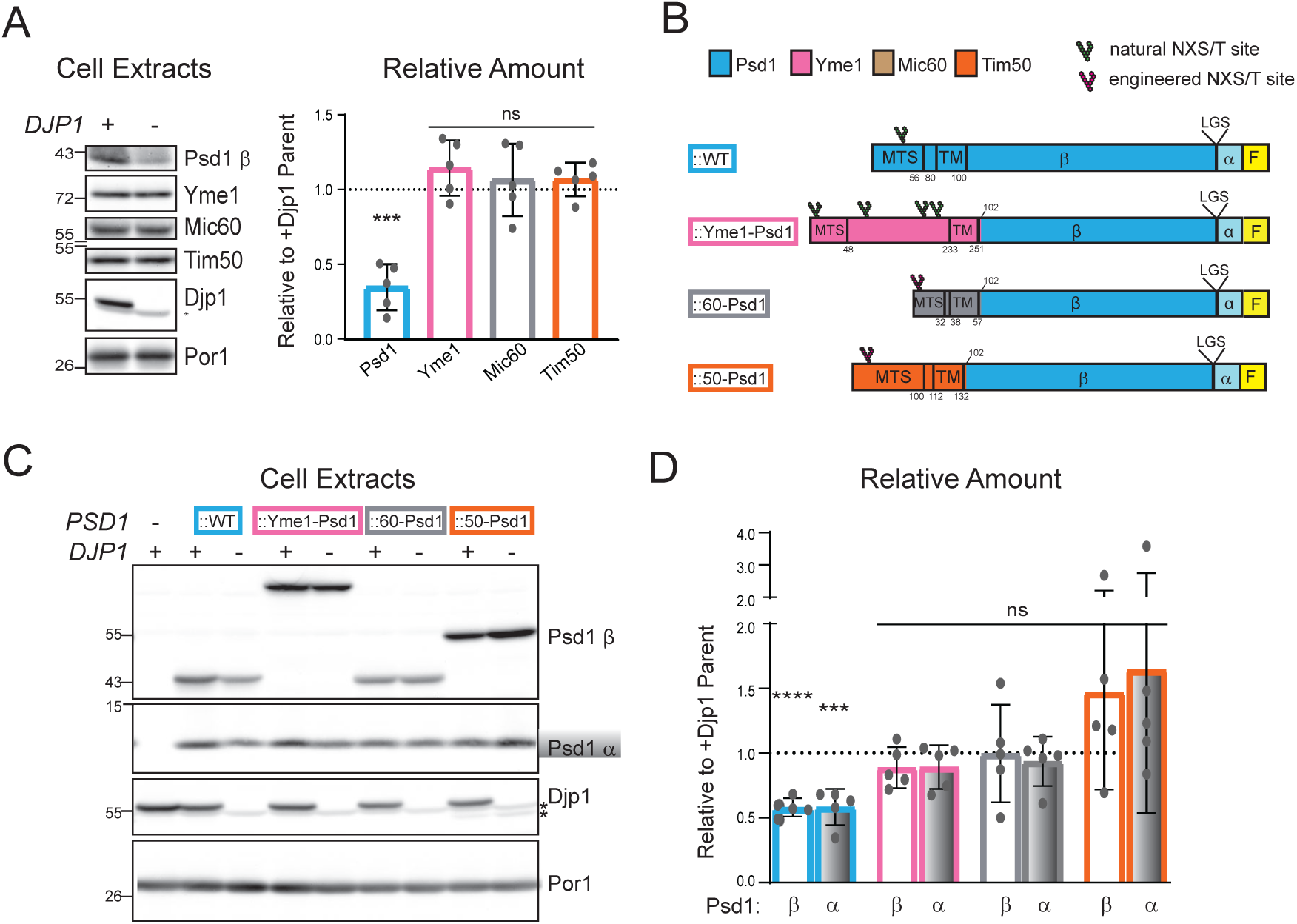
Djp1-dependence is contained within the N-terminal 102 amino acids of Psd1. **(A)** WT and *djp1*Δ cell extracts from yeast grown in rich lactate media were immunoblotted as indicated. Por1 acted as a loading control. The relative amounts of Psd1, Yme1, Mic60, and Tim50 in the absence versus the presence of Djp1; wildtype amounts were set to 1.0 (mean ± SD for n = 5 biological replicates). Significant differences (ns, not significant; 3 symbols, *P* ≤ 0.001) versus wildtype were determined by Welch’s t tests. **(B)** Schematic of N-terminal chimeric constructs. MTS (mitochondrial targeting signal) and TM (transmembrane) domain residues are listed. Psd1 β and α subunits are delineated by the LGS motif. All constructs have a C-terminal 3XFLAG tag. **(C)** Cell extracts from Djp1 containing or lacking *psd2*Δ*psd1*Δ yeast transformed with the indicated chimera were immunoblotted as listed. Por1 acted as a loading control. **(D)** The Por1-normalized amount of Psd1 β and α subunit from the designated chimera were determined relative to their respective Djp1-containing parent (mean ± SD for n = 5 biological replicates). Significant differences (ns, not significant; 3 symbols, *P* ≤ 0.001; 4 symbols, *P* ≤ 0.0001) versus the respective Djp1-containing parent were determined by unpaired t tests.

To further refine the location of the Djp1-sensitivity motif in the first 100 amino acids of Psd1, we generated a second set of Tim50-Psd1 chimeras (Fig 5A). The MTS of Psd1 is removed upon its import into the IM by the sequential action of two matrix localized proteases, the matrix processing peptidase and Oct1 [29]. The mature N-terminus of Psd1 post-import and cleavage is Gly57. The transmembrane domain of Psd1 spans from 80-100. Using a similar strategy in the context of Mic60-based fusions and EGFP tagging, it was previously concluded that the TM domain of Psd1 is necessary and sufficient for its localization to the ER [62]. However, in addition to its TM domain, the chimeric construct supporting this conclusion also included the preceding 23 amino acids upstream of the formal membrane spanning segment following MTS cleavage. As such, the second set of Tim50-Psd1 chimeras were designed to further distinguish between the TM domain alone (Psd1 TM) versus the TM domain and the upstream small stretch of residues post-MTS cleavage (Psd1 TM+). Like the original Tim50-Psd1 chimera (termed here Psd1 IMS), Psd1 TM and Psd1 TM+ were both catalytically active as seen by the appearance of free α subunit (Fig 5B). Perhaps contrary to expectations [62], the relative amount of each new chimera was unaltered when Djp1 was missing (Fig 5C), indicating that the MTS of Psd1 is necessary for its steady state accumulation to be controlled by Djp1.

**Fig 5.**
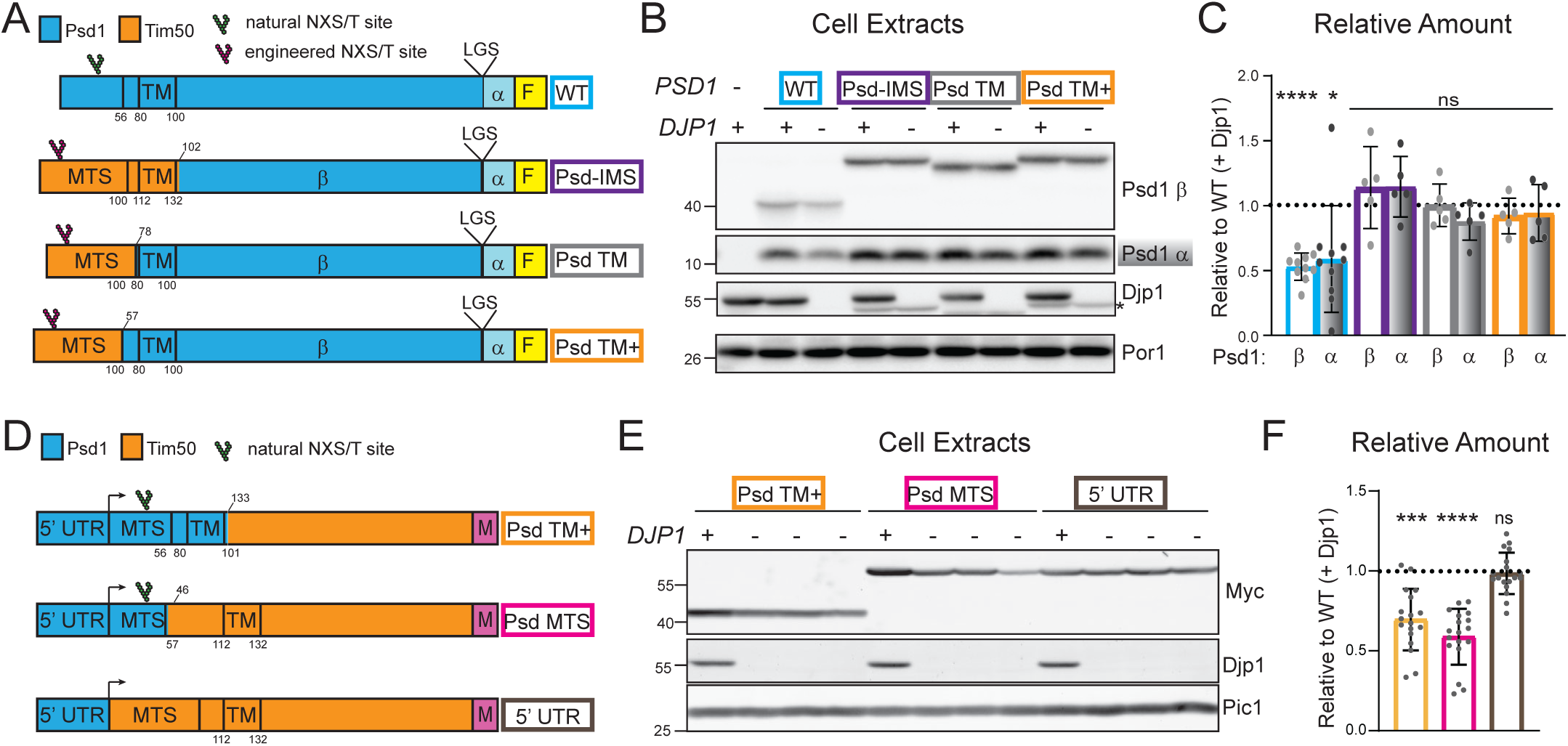
The MTS of Psd1 is necessary and sufficient for Djp1-dependence. **(A)** Cartoon of Tim50-Psd1 chimeric constructs; all constructs have a 3XFLAG tag at the C-terminus. **(B)** Cell extracts derived from the indicated strains grown at 30°C in rich dextrose were immunoblotted for Psd1 β, Psd1 α (FLAG), and Djp1; Por1 acted as loading control. **(C)** The Por1-normalized amount of Psd1 β and α subunit from the designated chimera were determined relative to their respective Djp1-containing parent (mean ± SD, n = 5-10). Statistical differences (ns, not significant; 1 symbol p<0.05; 4 symbols p<0.0001) for Psd1 β and α subunits compared to corresponding Djp1-containing parent were determined by unpaired t-test; ns, not significant. **(D)** Cartoon of Psd1-Tim50 chimeric constructs; all constructs have a 3XMyc tag at the C-terminus and are downstream of the *PSD1* 5’ UTR. **(E)** Cell extracts derived from the indicated strains grown as in (B) were immunoblotted for Myc and Djp1; Pic1 acted as loading control. Each lane for chimeras lacking Djp1 are from individual clones. **(F)** The Pic1-normalized amount of each Myc-detected chimera was determined relative to its Djp1-containing parent (mean ± SD, n = 6 total for Djp1 containing parents, 3 biological replicates of 2 parents per chimera; n = 18 total for Djp1-lacking daughters, 3 biological replicates of 3 daughters per parent). Statistical differences (ns, not significant; 3 symbols p < 0.001; 4 symbols p<0.0001) compared to corresponding Djp1-containing parent were determined by unpaired t-test.

To determine if the Psd1 MTS is sufficient to confer Djp1-sensitivity, we generated a final set of Psd1-Tim50 chimeras, each containing a C-terminal 3XMyc tag and placed downstream of the 5’UTR of *PSD1* (Fig 5D). Indeed, replacing the MTS of Tim50 with that of Psd1 resulted in a chimera (Psd1 MTS) whose relative amount was reduced in the absence of Djp1 (Fig 5E and 5F). Since the abundance of Tim50 under control of the *PSD1* 5’UTR was not Djp1-sensitive, this reduction is not likely to be transcriptionally based, consistent with our earlier finding (Fig 1C). Thus, the MTS of Psd1 is both necessary and sufficient to make a mitochondrial precursor sensitive to cellular Djp1 status.

### Only a few mitochondrial proteins are sensitive to cellular Djp1 status

Djp1 was originally assigned a specific role in peroxisomal protein import and maturation [80] and only more recently tied to the import and/or accumulation of a small cohort of mitochondrial precursors [41, 42, 61]. To establish a comprehensive list of Djp1-dependent mitochondrial clients, we isolated mitochondria from WT and *djp1*Δ yeast grown in rich dextrose or rich lactate media, confirmed that in the absence of Djp1, the Psd1 precursor did not accumulate in the ER-enriched P40 fraction and was reduced in mitochondria (Fig 6A and 6B), and then performed TMT proteomics on three individual mitochondrial preparations per genotype (Fig 6C and 6D). Our TMT proteomics data revealed changes in protein abundance of only a select number of proteins, most of which were residents of either peroxisomes (light blue) or mitochondria (red), and sometimes both (Fig 6C and 6D; purple). Only Psd1 and the known mitochondrial Djp1 clients, Mim1 and Mim2 [61], were significantly reduced in the absence of Djp1 regardless of metabolic state (Fig 6C-6E). Other mitochondrial proteins displayed the opposite pattern and were selectively increased in the absence of Djp1 in glycolytic dextrose (Mpc2, Mpc3, Cox23 and Cmc2) or respiratory lactate (Mtm1, Mfm1, Pad1, Cld1, and Sdh6) media (Fig 6E). Most peroxisomal proteins that were impacted by the absence of Djp1 were only reduced in respiratory conditions; only a handful (Aat2, Ant1 and Pmu1) were also decreased in dextrose-derived mitochondria. It is presently unclear if the changes in these peroxisomal proteins reflect their detection in peroxisomes that co-fractionate with mitochondria and/or alterations in their normal subcellular distribution when Djp1 is missing.

**Fig 6.**
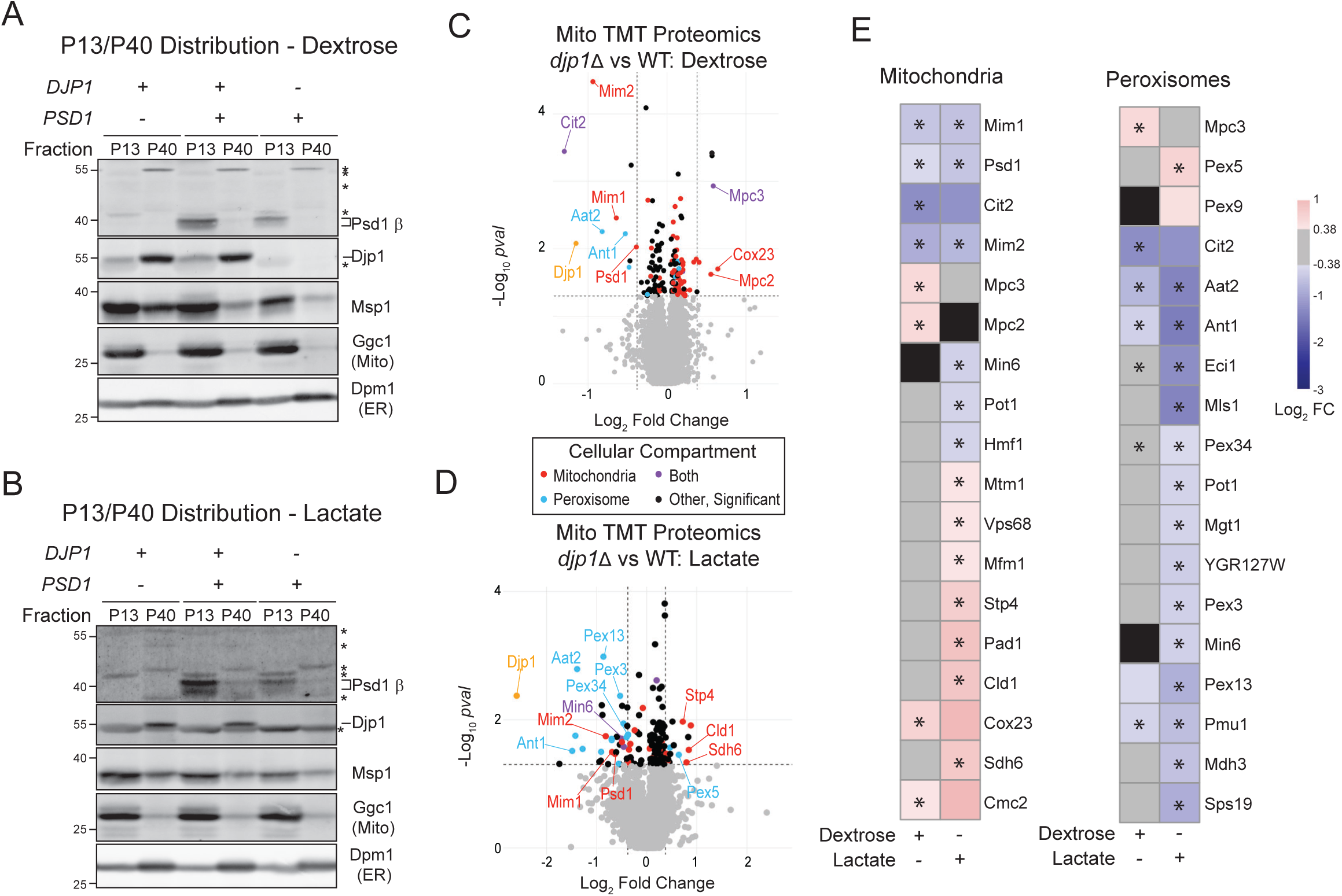
The amount of only a limited number of mitochondrial proteins are changed in the absence of Djp1. P13 (Mitochondria) and P40 (ER) fractions from yeast of indicated genotypes grown in **(A)** rich dextrose or **(B)** rich lactate were immunoblotted as indicated. **(C)** Tandem mass tag (TMT) comparison of mitochondrial proteomes from WT and *djp1*Δ yeast grown in rich dextrose (n = 3 preps each). **(D)** TMT comparison of mitochondrial proteomes from WT and *djp1*Δ yeast grown in rich lactate (n = 3 preps each). **(E)** Heatmaps of mitochondrial and peroxisomal proteins identified in TMT proteomics analysis of mitochondria isolated from WT and *djp1*Δ yeast grown in rich dextrose or lactate as indicated. Any protein that met the log2FC inclusion criteria for only one sample group was colored gray for the sample group in which the protein did not meet inclusion criteria and any protein that was not included in the dataset for one of the sample groups was colored black. Proteins that had log2FC in *djp1*Δ vs WT with p-value ≤0.05 are labeled (*).

Given the role of Djp1 in peroxisomal biogenesis (Fig 6C-6E and [80]) and an emerging link between mitochondria and peroxisomal biogenesis [81–83], we next tested the possibility that the steady state amount of Psd1 is sensitive to general peroxisomal defects. However, the steady state abundance of Psd1 was like WT in yeast lacking Pex3, Pex5, Pex7 or Pex13 (Fig 7A and 7B). This finding indicates that the roles of Djp1 in peroxisomal protein import and biogenesis and in influencing the amount of Psd1 in mitochondria are likely distinct.

**Fig 7.**
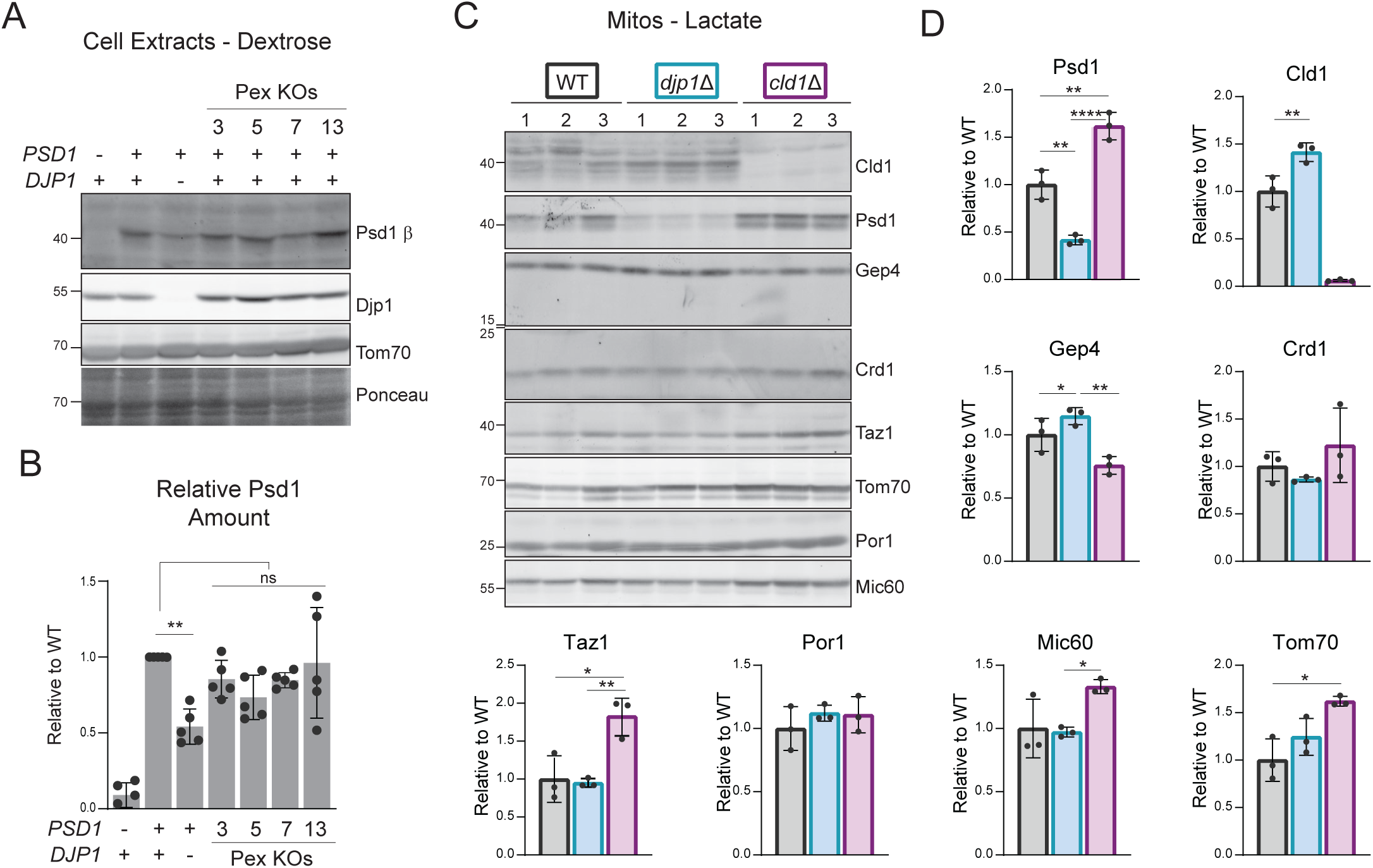
Mitochondrial phospholipid biosynthetic proteins are not generally sensitive to Djp1 absence. **(A)** The Psd1 β subunit was detected by immunoblot in yeast cell extracts of the indicated genotype grown in rich dextrose. Tom70 and total protein stain acted as loading controls. **(B)** The relative amount of Psd1 versus wildtype, normalized to total protein stain, was determined (mean ± SD for n =5 biological replicates). Significant differences (ns, not significant; 2 symbols, *P* ≤ 0.01) versus wildtype were determined by one-way ANOVA with Tukey’s multiple comparisons. **(C)** Mitochondria from lactate-grown yeast of indicated genotype were harvested by differential centrifugation and equal protein amounts resolved by SDS-PAGE and immunoblotted as listed. Each lane is a distinct mitochondrial preparation for each genotype. **(D)** The relative amount of each listed protein versus wildtype was determined (mean ± SD for n =3 biological replicates). Significant differences (1 symbol, *P* ≤ 0.05; 2 symbols, *P* ≤ 0.01; 4 symbols, *P* ≤ 0.0001) versus wildtype were determined by one-way ANOVA with Tukey’s multiple comparisons.

Of the few changes in mitochondrial protein amounts detected in the absence of Djp1, Cld1 was of immediate interest since it, like Psd1, functions in mitochondrial phospholipid metabolism. Cld1 deacylates newly synthesized CL, a four-acyl chain containing phospholipid exclusively made in mitochondria, which is then reacylated by the transacylase, Taz1 [84, 85]. In so doing, Cld1 plays an essential role in establishing the final acyl chain character of cardiolipin [84]. To test the possibility that Djp1 has a broader role in mitochondrial phospholipid metabolism, we determined the abundance of various proteins in WT, *djp1*Δ, and *cld1*Δ mitochondria isolated from rich lactate cultures (Fig 7C). Importantly, we confirmed by immunoblot the distinct patterns of Psd1 (decreased) and Cld1 (increased) when Djp1 is missing, and additionally determined that the PG-phosphate phosphatase, Gep4 [86], was also increased in the absence of Djp1 (Fig 7D). The steady state amounts of CL synthase, Crd1, and Taz1 were like WT in *djp1*Δ mitochondria. Unexpectedly, the protein levels of Psd1 and Taz1 were increased in *cld1*Δ mitochondria, which could reflect crosstalk between the PE and CL biosynthetic pathways for Psd1 and/or feedback control since Cld1 functions immediately upstream of Taz1. From these results, we conclude that Djp1 does not have a general regulatory role related to mitochondrial phospholipid metabolism. Further, the variable nature of the impact of Djp1 loss on the steady state amounts of select mitochondrial proteins suggests that it likely modulates mitochondrial homeostasis by more than a single, ER-SURF related mechanism.

### Djp1 has Psd1 and non-Psd1 related roles in mitochondrial phospholipid metabolism

In the absence of Djp1, only three mitochondrial proteins ‒ Psd1 and the OMM insertases Mim1 and Mim2 ‒ showed reduced steady state amounts in a metabolic state agnostic manner. Therefore, we performed a Psd-centric epistasis analysis to determine if perhaps Djp1 maintains cellular PE levels when the cellular redundancy in PE biosynthetic pathways in reduced (Fig 8). While Psd1 is the major source of cellular PE in yeast [22], there are three additional non-mitochondrial processes that can contribute to PE production in yeast. Psd1 and Psd2 are each sufficient for growth on synthetic defined media lacking ethanolamine, whereas yeast lacking both Psds are ethanolamine or lyso-PE auxotrophs [20, 33]. The absence of Djp1 resulted in a ∼50% drop in Psd1 amounts regardless of Psd2 status (Fig 8A). Despite this significantly decreased Psd1 abundance, yeast lacking both Psd2 and Djp1 grew just as well as their Djp1-containing parents on both ethanolamine-free glycolytic (SD, synthetic defined with dextrose) and respiratory (SDLac, synthetic defined with lactate) media (Fig 8B and 8C). Given that in these conditions the CDP-ethanolamine pathway is inactive due to a lack of ethanolamine and Psd2 is missing, these results argue that Djp1 does not act to safeguard cellular PE metabolism.

**Fig 8.**
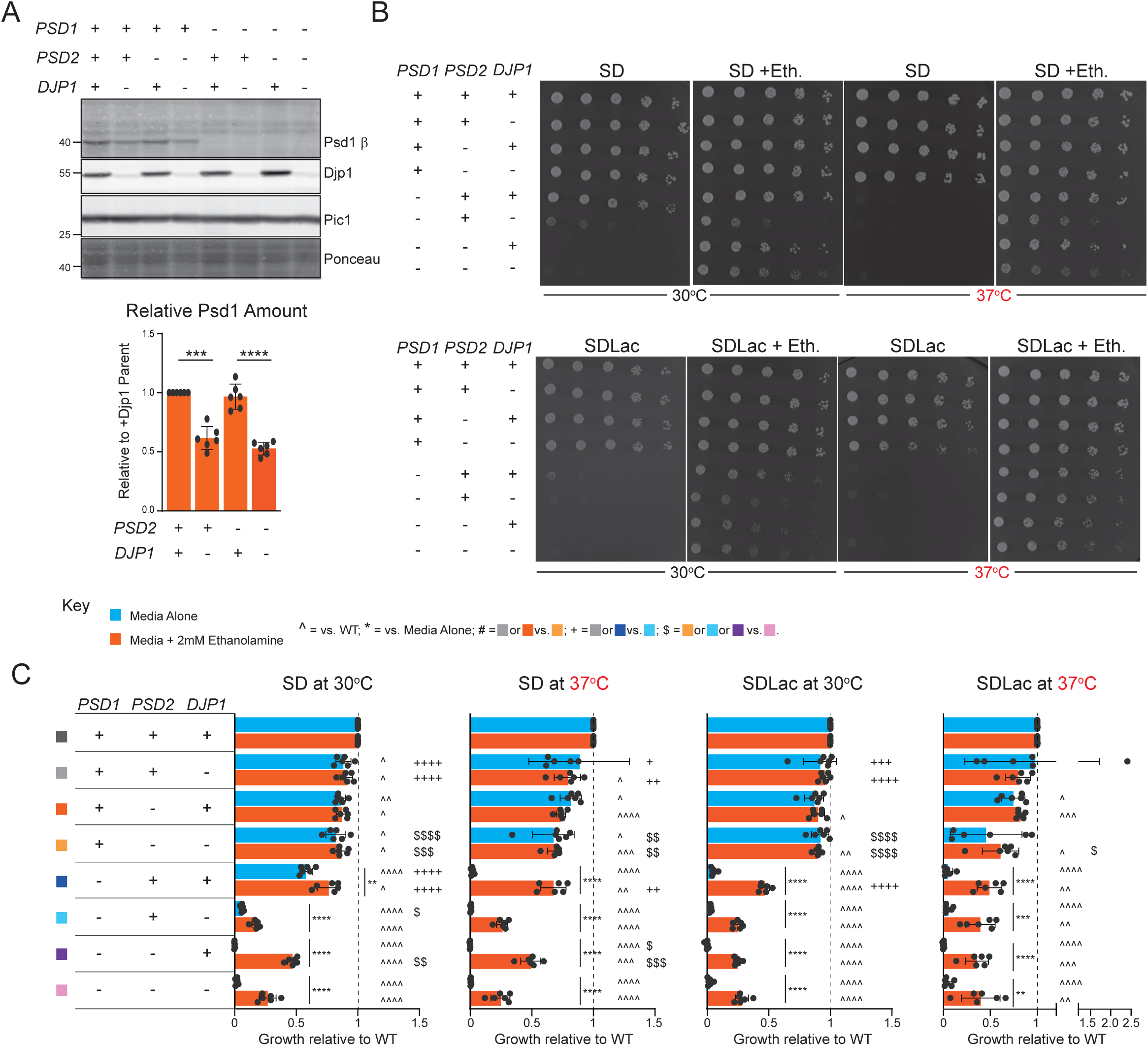
Combined loss of Psd1 and Djp1 results in a synthetic sick phenotype on dextrose and decreases ethanolamine-based rescue. **(A)** Cell extracts derived from the indicated strains grown at 30°C in rich lactate were immunoblotted for Psd1 β and Djp1; Pic1 and total protein stain acted as loading controls. The relative Pic1-normalized amount of Psd1 was determined with the protein levels of corresponding parent strains that express Djp1 set to 1.0 (mean ± SD for n = 6 biological replicates). Statistical differences (3 symbols p < 0.001; 4 symbols p<0.0001) compared to corresponding Djp1-containing parent were determined by Welch’s t-test. **(B)** The indicated strains were spotted onto SD and SDLac plates, each with or without 2mM ethanolamine, and grown for 2 (SD) and 3 (SDLac) days at 30°C or 3 (SD) and 5 (SDLac) days at 37°C. **(C)** Growth of each strain genotype relative to wildtype yeast was determined (mean ± SD for *n* = 6 biological replicates). Statistical differences compared to wildtype (^) between *psd2*Δ*djp1*Δ (#) and *psd1*Δ*djp1*Δ (+) and their two respective single deletion parents, between *psd2*Δ*psd1*Δ*djp1*Δ its three double deletion parents ($) were determined by one-way ANOVA with Tukey’s pairwise comparisons; differences as a function of ethanolamine (*) were calculated by unpaired student t tests.

Interestingly, the combined absence of Psd1 and Djp1 resulted in a synthetic sick growth defect on SD at 30℃, suggesting that Psd1 and Djp1 function in parallel pathways, whereas on SD at 37℃ or SDLac at both temperatures, Psd1 was epistatic to Djp1 (Fig 8B and 8C). The growth-boosting benefit provided by supplemental ethanolamine was significantly reduced in yeast lacking Psd1 and Djp1 (SD and SDLac) or Psd1, Psd2 and Djp1 (SD only) compared to their respective Djp1-containing parents. This latter finding suggests that Djp1 may facilitate the ability of ER-synthesized PE to rescue the loss of its normal production in the IMM.

Finally, we sought to directly determine how the loss of Djp1 impacts PE metabolism in the absence of Psd1 and/or Psd2 (Fig 9). Consistent with their nonexistent growth phenotype, the levels of PE, its precursor, PS, and its product, phosphatidylcholine (PC), were largely unchanged in *psd2*Δ, *djp1*Δ, and *psd2*Δ*djp1*Δ versus WT yeast in both SD (Fig 9A and 9B) and SDLac (Fig 9 C and D). In SD media, PE was decreased and PS and PC increased upon the combined loss of Psd1 and Djp1 relative to single *psd1*Δ and *djp1* strains; these changes resulted in reduced PE:PS and PE:CL and elevated PC:PE ratios (Fig 9A and 9B). In SDLac, the PE and PC levels and PE:PS, PC:PE, and PE:CL ratios were the same in yeast lacking Psd1 or both Psd1 and Djp1, but different than single *djp1*Δ yeast (Fig 9C and 9D). These results further underscore that the genetic relationship between Psd1 and Djp1 differs based on whether yeast are allowed to produce energy via glycolysis (SD; Psd1 and Djp1 function in parallel pathways) or forced to do so via oxidative phosphorylation (SDLac; Psd1 is epistatic to Djp1). In SD media, PE was not increased by the inclusion of ethanolamine for any tested genotype (Fig 9A and 9B) whereas it was when the same strains were grown in SDLac (Fig 9C and 9D). This indicates that the CDP-ethanolamine Kennedy pathway and/or the ability of PE made by it to gain access to mitochondria is sensitive to cellular metabolic state. Interestingly, PS was significantly increased by ethanolamine in *psd1*Δ*djp1*Δ yeast grown in SD (Fig 9A and 9B) but not SDLac (Fig 9C and 9D). PS levels were also high in *psd1*Δ*psd2*Δ and *psd1*Δ*psd2*Δ*djp1*Δ yeast grown in ethanolamine-spiked SD, similar to *psd1*Δ*djp1*Δ, whereas it was like WT in yeast lacking Psd2 and Djp1. In SDLac with ethanolamine, PS was elevated in yeast lacking Psd1, Psd2, and Djp1 compared to any double knockout parent. Phosphatidylinositol (PI) was reduced by ethanolamine in yeast lacking Psd1 alone or with Djp1 in both SD and SDLac media; however, in the context of Psd1-lacking yeast, ethanolamine had an additive effect with the additional loss of Djp1 in SD, but not SDLac (Fig 9). Finally, PI levels were the same in *psd2*Δ*psd1*Δ and *psd2*Δ*psd1*Δ*djp1*Δ yeast grown in SD with ethanolamine (Fig 9A and 9B) but reduced in the latter strain in ethanolamine-containing SDLac (Fig 9C and 9D). From these combined results, we conclude that Djp1 does not function as a cellular PE metabolism safeguard, at least in the conditions tested, but does have unanticipated roles in mitochondrial phospholipid metabolism that interact with Psd1 in a metabolic state sensitive manner.

**Fig 9.**
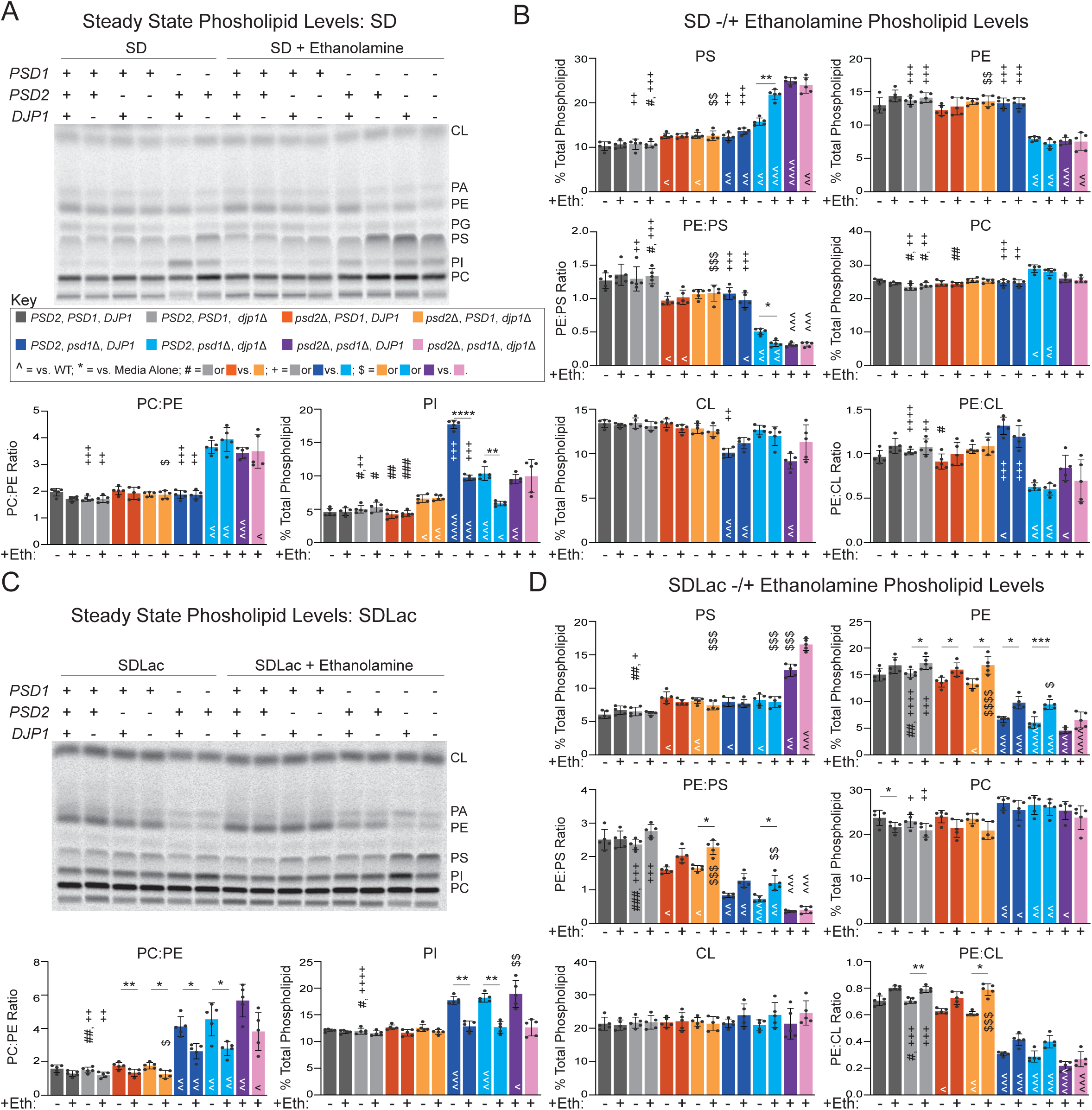
Combined loss of Psd1 and Djp1 broadly perturbs mitochondrial phospholipid levels in a metabolically-sensitive manner. Mitochondrial phospholipids were labeled overnight with ^14^C-acetate in the indicated yeast grown in SD **(A)** or **(C)** SDLac, each with or without 2mM ethanolamine, extracted, and separated by TLC. **(B, D)** Quantitation of the listed mitochondrial phospholipid levels or ratios (mean ± SD for *n* = 6 biological replicates). Significant differences compared to wildtype (^), between *psd2*Δ*djp1*Δ (#) and *psd1*Δ*djp1*Δ (+) and their two respective single deletion parents, between *psd2*Δ*psd1*Δ*djp1*Δ its three double deletion parents ($), and as a function of ethanolamine (*) were determined by one-way ANOVA with Tukey’s pairwise comparisons.

## Discussion

In this study, we set out to test the hypothesis that the mitochondrial PE synthase and IMM resident, Psd1, is a novel substrate of the recently appreciated Djp1-dependent ER-SURF pathway. Consistent with this model, we determined that the ER-associated Hsp40 cochaperone Djp1 was required for the full accumulation of Psd1 in mitochondria following growth in either glycolytic or respiratory conditions via a post-transcriptional mechanism. Having identified Psd1 as a likely biogenic client of Djp1, we then aimed to use this knowledge to define molecular mechanisms underlying Djp1-dependent regulation of mitochondrial proteins and interrogate its role in sustaining overall mitochondrial function.

Psd1 dependency on Djp1 is highly specific and unshared amongst other Hsp40 family members tested. Here, we demonstrated that Psd1 steady state levels were unaffected and not rescued by the loss or overexpression of additional Hsp40 family members previously implicated in mitochondrial precursor targeting and import (Ydj1, Xdj1, and Sis1 [66, 67, 87, 88]), assigned as partially redundant with Djp1, albeit in non-mitochondrial contexts (Ydj1 and Caj1 [68, 69]), or associated with the ER like Djp1 (Erj5 [70] and Hji1 [71]). Further, the ability to fully support the steady state load of Psd1 was dependent on each known Djp1 domain and required its Hsp70-stimulating activity.

Like Psd1, Djp1 is also important for the import and steady state accumulation of Mim1 and Mim2 in a manner that is unique amongst cytosolic Hsp40 family members [61]. Our proteomics analyses confirmed that Mim1 and Mim2 amounts are reduced in the absence of Djp1 and further demonstrated that this relationship is also insensitive to cellular metabolic state. In fact, Mim1, Mim2, and Psd1 were the only mitochondrial proteins that were decreased in Djp1-null yeast in both glycolytic and respiratory growth conditions (Fig 7). Unlike Psd1, Mim1 and Mim2 are OMM proteins that form the MIM insertase, a complex that promotes the import and membrane integration of OMM proteins with α-helical TM domains [4, 5, 89]. While Psd1, Mim1, and Mim2 all contain a single TM domain, their membrane topology in the IMM or OMM are flipped [27, 90]. Interestingly, the TM domain of Mim1 can functionally rescue *mim1*Δ yeast when overexpressed, indicating that the N- and C-termini of Mim1 are dispensible for OMM targeting and MIM insertase function [90]. While Djp1 binds the Mim1-TM domain [61], in addition to other peptide stretches, the minimal Djp1-sensititivity motif in Mim1 and Mim2 are presently unresolved.

Based on the comparative proteomics of WT and *djp1*Δ mitochondria, the majority of mitochondrial membrane proteins regardless of submitochondrial localization do not share the post-transcriptional dependency of Psd1 on Djp1. Factors that allow for the selective recognition of a small subset of proteins are presently unknown. The MTS has emerged in recent years as an active participant in protein import and mitochondrial stress signaling [91–94]. While MTSs do not possess a single consensus sequence, they tend to form positively charged amphiphilic α-helices that are 15-60 residues long [40]. MTSs are now appreciated to contain a plethora of nuanced information that impact various import parameters, such as import efficiency, rate, lag-time, and duration, all of which can modulate mitochondrial function and initiate stress responses critical for cellular fitness [92]. Our demonstration that the Psd1 MTS is both necessary and sufficient to confer Djp1 dependency indicates that the Psd1 MTS possesses an additional layer of information not shared by most MTS-containing mitochondrial proteins. Interestingly, Mim1 and Mim2 lack an MTS and rely on non-cleavable internal sequences for targeting. Thus, it appears likely that Djp1 engages its mitochondrial clients by multiple means. In the case of Psd1 and Djp1, it must be noted that we have not established that they physically interact, leaving open the possibility that the role of Djp1 as a determinant of Psd1 amounts is indirect. Further, the chimeras employed to test for the sufficiency of the Psd1 MTS all contained a TM domain just downstream (Fig 5D-5F). Thus, how Djp1 detects the MTS of Psd1, whether this involves another targeting factor, and if a juxtaposed TM domain is also a critical determinant are open questions for future pursuit.

Our understanding of how different mitochondrial protein targeting pathways coordinate or work separately is limited. Previous work has shown that Djp1 works downstream of the GET pathway to mediate the import of Oac1 [43]. We demonstrated that Psd1 steady state levels remained unchanged in the absence of Get3 suggesting that cooperation of the GET and ER-SURF pathways is critical to only a subset of mitochondrial proteins. We also failed to invoke a role for SRP-dependent targeting as a determinant of Psd1 accumulation, but did identify an unexpected consequence of a defective SRP-independent pathway: Psd1 amounts were significantly increased. It was recently shown that Sec72 deletion has substrate-dependent effects on protein secretion, including enhanced secretion of proteins with strongly hydrophobic signal peptides [95]. Thus, the increase in Psd1 abundance in the absence of Sec72 may reflect a client-specific consequence of Sec72 loss, although whether this results from altered synthesis, targeting, or processing remains unknown.

Unlike previous reports for Mim1, Oxa1, or Oac1 [41, 43, 61], we failed to detect an accumulation of Psd1 on the ER upon loss of Djp1. We suggest two possibilities for this difference: 1) Psd1 dependency on Djp1 is separate from Djp1’s role in ER-SURF; or 2) Psd1 was undetectable at the ER due to its rapid detection and turnover by quality control mechanisms therein. Recent work has suggested that the relative hydrophobicity of internal protein segments, e.g. TM domains, in integral IMM proteins dictates their propensity for ER targeting and need for ER-SURF [42]. The TM domains of Psd1 (Djp1-dependent) and Tim50 (Djp1-independent) both rank as strongly hydrophobic with GRAVY (grand average of hydropathy) scores of 1.3 and 0.93, respectively. The importance of the TM domain and its relative hydrophobicity in conferring Djp1 dependency awaits systematic evaluation. We recently demonstrated that non-imported, nonfunctional Psd1 is partially directed to the ER for subsequent degradation by the ubiquitin-proteasome system [64]. As such, it is possible that the failure to detect ER accumulation of Psd1 in the absence of Djp1 reflects its swift resolution in this compartment. Whether loss of Ema19 [48] and/or Spf1 [50], each implicated in removing proteins mistargeted to the ER, or chemical perturbation of the ubiquitin-proteasome system would result in ER accumulation of Psd1 when Djp1 is gone is an important follow up that would help understand variability in the fate of non-imported mitochondrial proteins. Djp1 is implicated in the import and steady state levels of a small, but diverse set of mitochondrial proteins; whether turnover of these substrates varies is unknown.

Djp1 is an established regulator of peroxisomal protein import and biogenesis [68, 80]. Consistently, our proteomics showed a reduction in the abundance of several peroxisomal proteins in the absence of Djp1. This observation raised the possibility that altered peroxisomal function may disrupt Psd1 steady state levels. Contrary to this possibility, we show that loss of several peroxisomal proteins failed to affect Psd1 steady state amounts as observed in absence of Djp1. Further, loss or overexpression of Caj1, another J-protein that exhibits functional redundancy with Djp1 in regard to peroxisomal biogenesis [68], failed to affect Psd1 steady state levels. When taken together, our findings support the idea that Djp1’s role in peroxisomal and mitochondrial biogenesis are distinct. The surface of the ER contains functional microdomains that regulates organelle biogenesis [96]. If and how pools of Djp1 are separated on the surface of the ER to carry out distinct biogenic functions remains to be elucidated.

Finally, we performed a cellular PE-focused epistasis analysis which uncovered a multifaceted involvement of Djp1 in mitochondrial phospholipid metabolism that is sensitive to the cellular metabolic state. In both their growth properties and phospholipid profiles, we determined that the genetic relationship between Psd1 and Djp1 was different when yeast were allowed to produce energy via glycolysis (Psd1 and Djp1 function in parallel pathways) or forced to do so via oxidative phosphorylation (Psd1 is epistatic to Djp1). Interestingly, PE and PI levels were significantly reduced upon the additional loss of Djp1 in Psd1-null yeast grown in glycolytic, but not respiratory, conditions (Fig 9). This suggests that Djp1 promotes mitochondrial PE and PI levels independent of Psd1 when glycolysis is the main source of energy. While extra-mitochondrially made PE can access this organelle, it has a limited ability to recover PE-associated functions normally provided by IM-produced PE [12, 35, 37, 97]. PI is made in the ER by Pis1 [98] and is a major component of mitochondrial membranes in yeast. How PE and PI move from the ER to mitochondria is unknown, but is likely mediated in part by ERMES [99–101], a protein complex that physically bridges ER and mitochondrial membranes [102] and is modeled to be most important in respiratory conditions [103]. How Djp1 promotes the mitochondrial uptake of PE, PI and perhaps other phospholipids when yeast are supported by glycolysis is an important ongoing extension of this unexpected discovery.

Factors such as Djp1 and the surface of the ER have emerged as critical contributors of mitochondrial protein targeting, but insights into substrate specificity, pathway redundancy, and impacts on broader mitochondrial function have been limited. Altogether, our study provides novel mechanistic insight into the regulation of the mitochondrial PE synthase, Psd1 by Djp1 and highlights the importance of the Psd1 MTS for this regulation. Moreover, we unveiled that Djp1 promotes normal mitochondrial phospholipid metabolism by metabolically-sensitive Psd1-dependent and independent mechanisms.

## Data and Code Availability

The accession number and DOI for the TMT-based proteomics data reported in this paper and deposited to the ProteomeXchange Consortium (http://proteomecentral.proteomexchange.org/cgi/GetDataset) are PRIDE: PXD081211 and 10.6019/PXD081211, respectively.

## Materials and Methods

### Molecular Biology

*DJP1*, along with its 5’ (533bp) and 3’ (336bp) untranslated regions (UTRs), was amplified from GA74-1A genomic DNA and cloned into the yeast centromeric plasmid, pRS316, or the yeast 2μ plasmid, pRS426, using convenient, endogenous BamHI and XbaI restriction sites. To generate a C-terminally tagged Djp1 construct, the SpyCatcher002 (SC) sequence was amplified from pDEST14-SpyCatcher (a gift from Mark Howarth (Addgene plasmid # 102827; http://n2t.net/addgene:102827; RRID:Addgene_102827) [104] and fused to the C-terminus of Djp1 in the pRS316-Djp1 plasmid by overlap-extension PCR. An 8x His tag was subsequently added to the 3’ end of the SC sequence by incorporating the tag in PCR primers, followed by a second round of overlap extension PCR. The Djp1 truncations (Djp1-J domain: 1^64-432aa; Djp1-J domain: 1^64-432aa; Djp1-F/G domain: 1-63^103-432; and Djp1-CTD: 1-103aa) and H34Q mutant, each driven by the native *DJP1* promoter (533bp 5’UTR) and subcloned into pRS316, were generated by overlap extension PCR using pRS316Djp1-SpyCatcher002-8xHis as the template. To express different SpyCatcher002-8xHis tagged yeast Hsp40 co-chaperones under the same regulatory control elements, yeast pRS316 was modified to contain the *AAC2* promoter (361 bps 5’UTR) and terminator (485 bps 3’UTR) sequences separated by a polylinker, generating pWizz. The ORFs for the Hsp40 co-chaperones Caj1 (type II), Sis1 (type II), Hlj1 (type II), Xdj1 (type I), Ydj1 (type I), and Erj5 (type III), representing different Hsp40 classes, were PCR amplified from GA74-1A genomic DNA, fused to SpyCatcher002-8xHis tag via overlap-extension PCR, and cloned into pWizz. IM-directed Psd1 chimeras in which the mitochondrial targeting signal (MTS) and transmembrane domain of Psd1 (1-101) was replaced by the equivalents from Tim50 (1-132; referred to in Figure 6 A,B as Psd1-IMS), Mic60 (1-57) or Yme1 (1-251), which all have the same IMM topology as Psd1 but are not Djp1 clients, have been described [64]. Additional Tim50-based chimeras were similarly produced by overlap extension PCR and resulted in Psd1 TM (1-100 of Tim50 attached to 78-3XFLAG tag of Psd1-3XFLAG) and Psd1 TM+ (1-100 of Tim50 attached to 57-3XFLAG tag of Psd1-3XFLAG). All chimeras containing the Tim50 MTS were also Q30N-modified by overlap extension to introduce an N-glycosylation signal (NSX) to detect potential targeting into the secretory pathway. To determine if the MTS of Psd1 was sufficient to confer Djp1-dependence, overlap extension PCR was used to generate Psd TM+ (5’ UTR and residues 1-101 of Psd1 fused to residues 133- end of Tim50), Psd MTS (5’ UTR and residues 1-57 of Psd1 fused to residues 46-end of Tim50), and 5’ UTR (5’ UTR of Psd1 fused to residues 1-end of Tim50). All three constructs contained the 3’ UTR of Tim50 (409 bps immediately downstream of stop codon) and were cloned into pRS305. Overlap extension PCR was then utilized to attach a 3XMyc tag onto the COOH terminus of Tim50 in each construct. All constructs generated for this study were verified by Sanger sequencing.

### Yeast strains and growth conditions

All yeast strains used in this study are listed in Table 1. Yeast expressing Psd1 with a 3XFLAG tag added to the C-terminus had Psd1-3XFLAG integrated into the *LEU2* locus [63] or instead knocked into the endogenous *PSD1* locus [25]. All pRS305-based Psd1 chimeras were linearized with AflII and integrated into the LEU2 locus of the *psd1*Δ*psd2*Δ (Psd1 chimeras placed downstream of N-terminal portions of other IM proteins) or wild type GA74-1A (Tim50 placed downstream of different N-terminal portions of Psd1) yeast strains. Clones were selected on synthetic complete dropout medium (0.17% (w/v) yeast nitrogen base (US Biological Y2035), 0.5% (w/v) ammonium sulfate, 0.2% (w/v) dropout mixture synthetic minus Leucine (US Biological D9526), 2% (w/v) dextrose, 2% (w/v) agar) -leucine (SC-Leu) and verified by immunoblot. Yeast transformed with non-integrating yeast plasmids (pRS316-, pRS426-, and pWizz-based) were selected and then maintained on synthetic complete lactate minus Uracil (SCLac-Ura; 0.17% (w/v) yeast nitrogen base, 0.5% (w/v) ammonium sulfate, 0.2% (w/v) dropout mixture synthetic minus Uracil (US Biological D9536), 0.05% (w/v) dextrose, 2% (v/v) lactic acid, 3.4mM CaCl_2_-2H_2_O, 8.5mM NaCl, 2.95mM MgCl_2_-6H_2_O, 7.35mM KH_2_PO_4_, 18.7mM NH_4_Cl, pH 5.5, 2% (w/v) agar) agar plates.

**TABLE 1.**
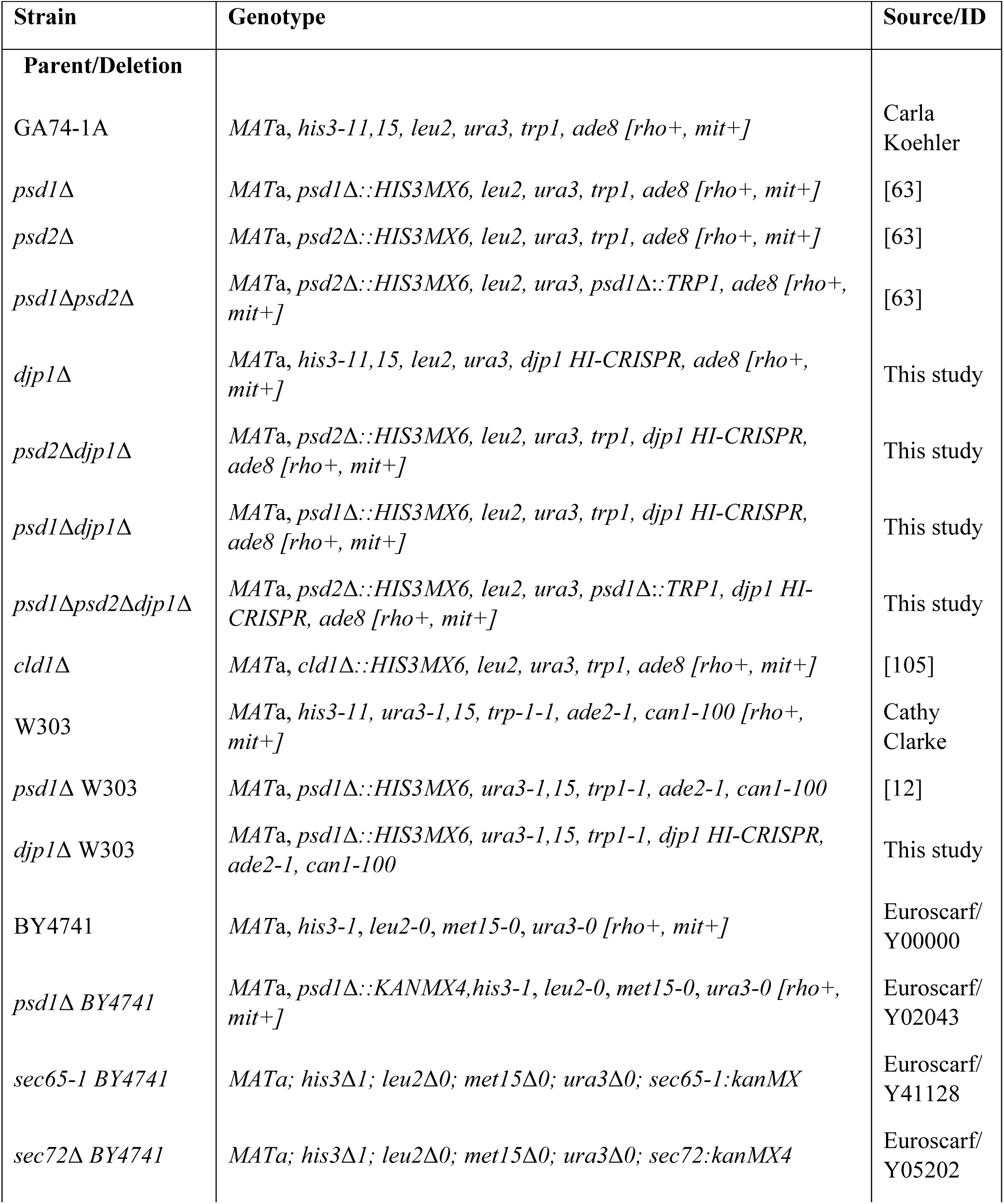
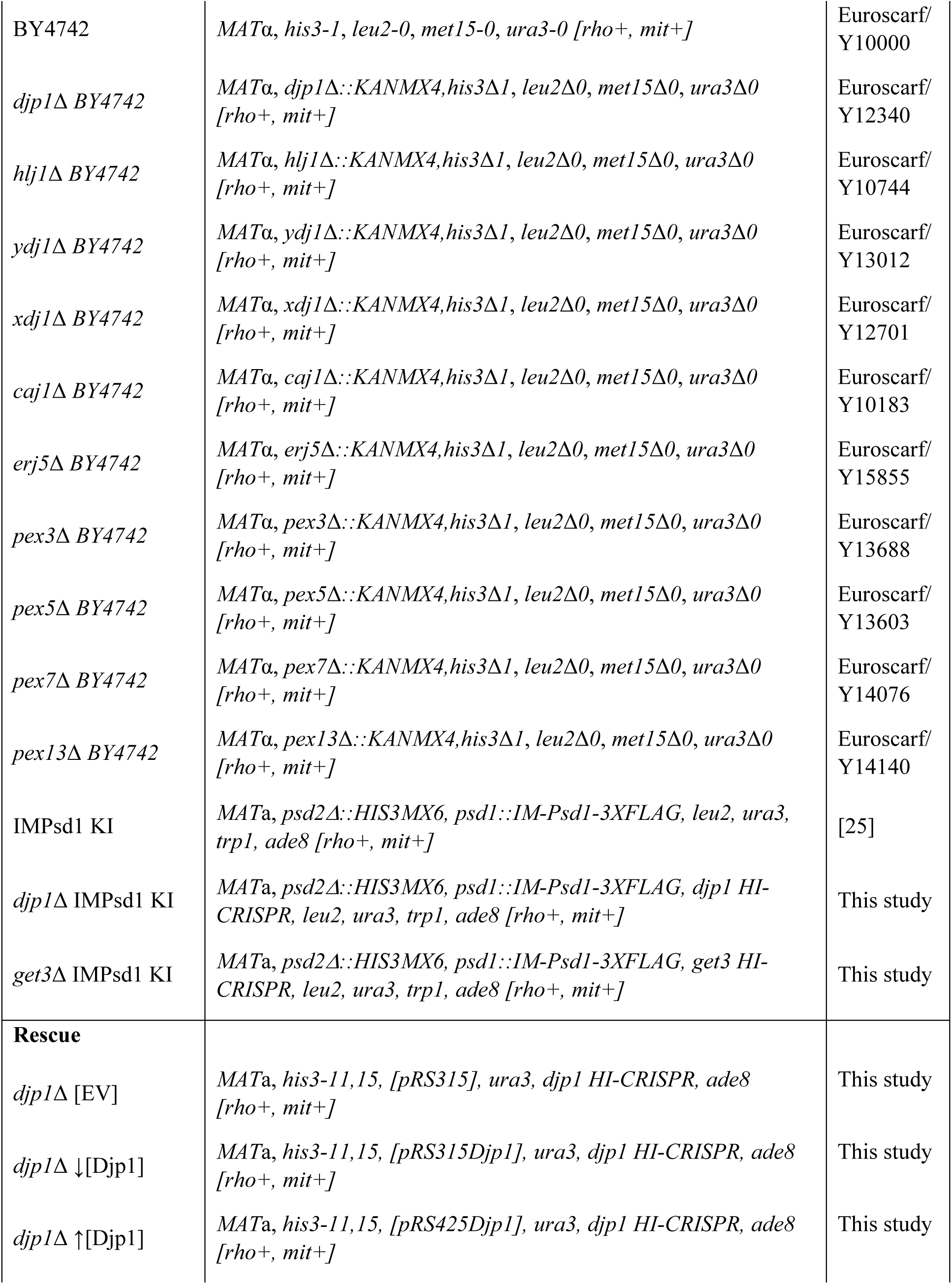
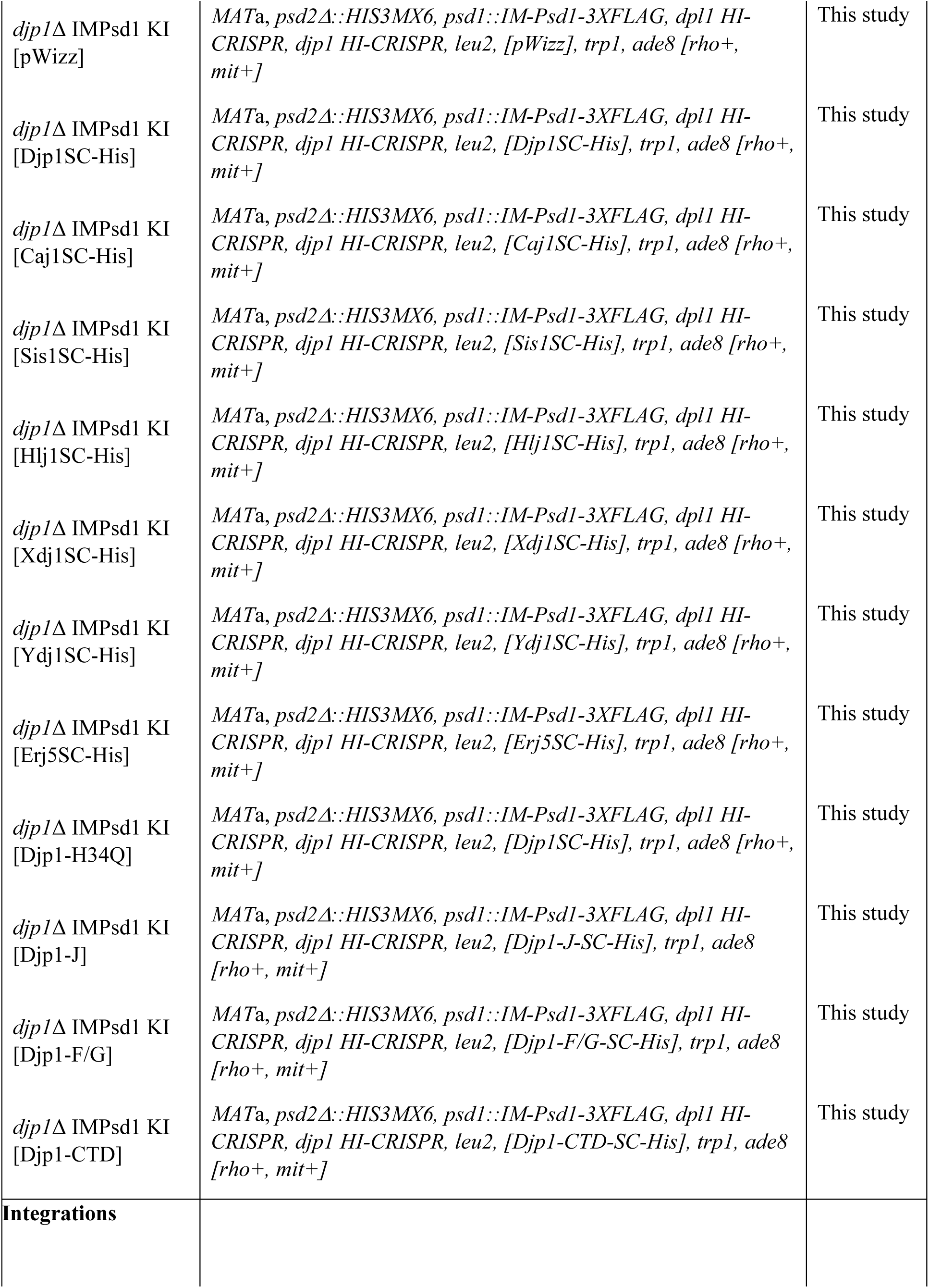
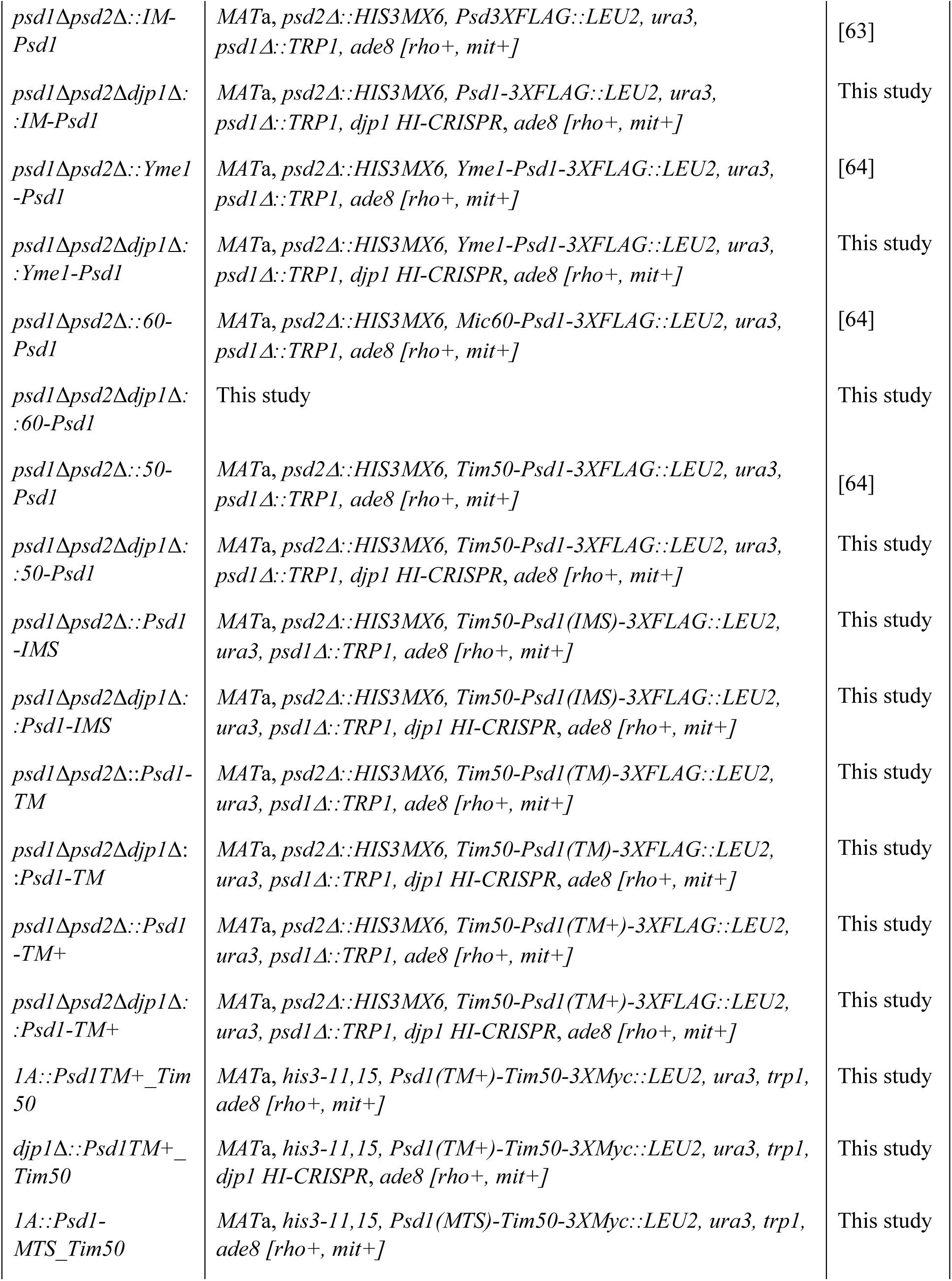
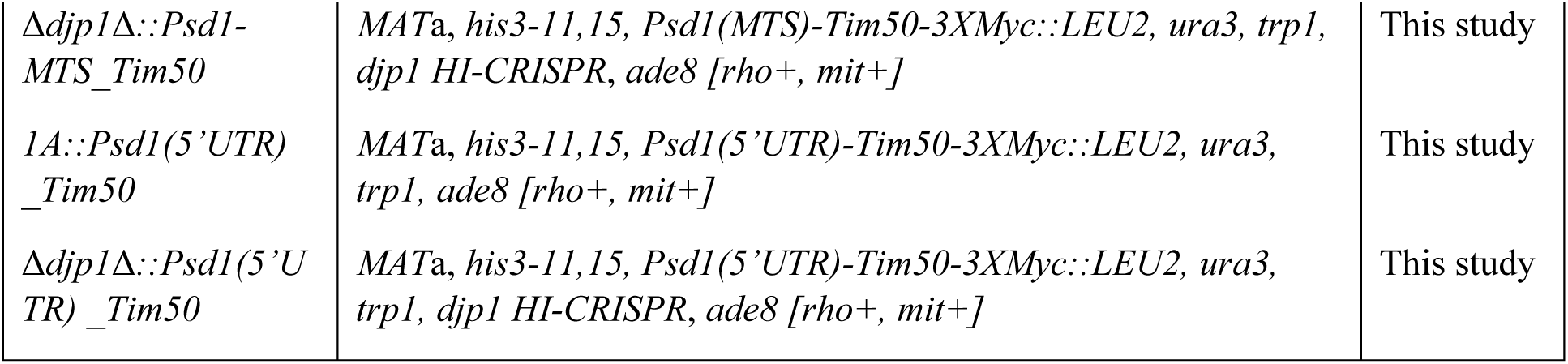
Strains used in this work.

Inactivation of the *DJP1* and *GET3* genes was achieved by homology-integrated clustered regulatory interspaced short palindromic repeats (CRISPR)-Cas (HI-CRISPR) [106]. CRISPR-Cas9 constructs were designed targeting *DJP1* or *GET3*, ordered as geneblocks (Integrated DNA Technologies) and assembled into the pCRCT plasmid (pCRCT was a gift from Huimin Zhao (Addgene plasmid #60621; http://n2t.net/addgene:60621; RRID:Addgene_60621), as described in [12, 31]. Both geneblocks included the 20bp CRISPR-Cas9 target and the homology repair template with 50bp homology arms on each side of the Cas9 recognition sequence. The homology repair template for the *DJP1* knockout replaced nucleotides 225-241 (17bp) with a 9bp segment while the *GET3*-directed homology repair template replaced nucleotides 601-618 with an 8bp segment (nucleotide numbers are with respect to the ORF start codon ATG sequence, defined as 1-3); both resulted in a premature stop codon and removed the PAM sequence to prevent re-cleavage by Cas9 once homology directed repair had occurred. The CRISPR-Cas9 system targeted the protospacer adjacent motif (PAM) sequence encoded by nucleotides 237-239 on the + strand of the ORF for *DJP1* and 615-617 on the + strand of the ORF for *GET3*. Successful targeting of these genes following transformation and selection was verified by immunoblotting.

Other than the yeast transformants with non-integrating plasmids (maintained on SCLac-Ura plates), yeast were maintained on either rich dextrose (YPD; 1% (w/v) yeast extract, 2% (w/v) tryptone, 2% (w/v) dextrose) or rich lactate plates (1% (w/v) yeast extract, 2% (w/v) tryptone, 0.05% (w/v) dextrose, 2% (v/v) lactic acid, 3.4mM CaCl_2_-2H_2_O, 8.5mM NaCl, 2.95mM MgCl_2_-6H_2_O, 7.35mM KH_2_PO_4_, 18.7mM NH_4_Cl, pH 5.5, 2% (w/v) agar). For spot-based growth studies, cultures were initially grown overnight in rich lactate media and used to generate a series of five 1:4 dilutions in sterile ddH_2_O starting with 4 x 10^-5^

OD_600_ cells/mL. An equal volume (3uL) of each dilution was spotted onto synthetic defined dextrose (SD; 0.67% (w/v) yeast nitrogen base with 0.5% (w/v) ammonium sulfate (RPI Y20040), 2% (v/v) complete amino acid mixture (added post-autoclave from a 50X filter-sterilized stock that contained 1g/L Adenine, 1g/L L-Arginine, 1g/L L-Histidine, 3g/L L-Leucine, 11.5g/L L-Lysine, 1g/L L-Methionine, 15g/L L-Threonine, 1g/L L-Tryptophan, 1g/L L-Uracil), 2% (w/v) dextrose, 2.5% (w/v) agar (Sigma 05038 or HiMedia RM301)) or synthetic defined lactate (SD-LAC; 0.67% (w/v) yeast nitrogen base with 0.5% (w/v) ammonium sulfate, 2% (v/v) complete amino acid mixture, 0.05% (w/v) dextrose, 2% (v/v) lactic acid, 3.4mM CaCl_2_-2H_2_O, 8.5mM NaCl, 2.95mM MgCl_2_-6H_2_O, 7.35mM KH_2_PO_4_, 18.7mM NH_4_Cl, pH 5.5, 2.5% (w/v) agar) agar plates, each +/- 2mM sterile-filtered ethanolamine hydrochloride, wrapped in parafilm and grown upside-down at either 30°C or 37°C. Yeast plates were imaged face-up without lids using a Chemidoc MP Imaging System (BioRad) and colorimetric acquisition mode after 2 or 3 days (30°C) or 3 or 5 days (37°C) of growth, as indicated in associated figure legend. The background subtracted volume (INT*mm) from two dilutions per genotype were averaged and growth relative to wild type yeast on the same plate determined.

### Preparation of yeast cell extracts

Other than the pRS316-based transformants, yeast were grown overnight in 2mL of either rich lactate (Figures 1B, 2B, 5, 6 and 7A) or rich dextrose (Figures 1A, 3 and 4A) media shaking at 220 rpm at 30°C; the pRS316-based transformants were grown overnight in 2mL SCLac-Ura. To test for a potential role of the SRP-dependent pathway, starter cultures of WT (BY4741) and *sec65-1* [73] yeast were grown overnight in 3mL of rich dextrose media at 23°C. The following day, 2 OD_600_ units of cells were transferred to new tubes, supplemented with fresh rich dextrose media to reach a final volume of 2mL at a density of 1 OD_600_/ml, and then incubated shaking at 220 rpm at 23°C or 35°C for 5 hours. After this period, 1 OD_600_ of cells were harvested in 1.5mL microcentrifuge tubes for 10 min at 845 × *g* at room temperature. Post-aspiration, cell pellets were resuspended with 1mL of water by a brief vortex after which 150 µL of a newly made NaOH/β-mercaptoethanol solution (1mL 2M NaOH: 80µL of β-mercaptoethanol) was added and the tubes inverted to mix before being placed on ice. Tubes were mixed every 2 min for 10 min before 75µL of 100% (w/v) trichloroacetic acid (TCA) was added to each tube. Once added, tubes were again mixed by inversion, placed on ice, and then incubated for another 10 min with regular mixing every 2 min. Proteins were collected with a 21,000 × *g* (room temperature or 4°C) centrifugation and isolated from the supernatant by aspiration. The protein pellets were washed with 1mL acetone to remove residual TCA and lipids, followed by a second 2 min centrifugation at 21,000 × *g*. For each sample, the acetone wash was decanted and the remaining pellet briefly dried on the bench before 30µL of 0.1M NaOH was added. After sitting at room temperature for 30-120min, protein samples were mixed with an equal volume of 2X reducing sample buffer (RSB), heated at 95°C for 5 minutes, and equal volumes (ranging from 6 to 8µL depending on target protein) of yeast extracts resolved by SDS-PAGE using custom-made 10-16% or 10-18% gradient gels for immunoblotting.

### Immunoblotting

Post-resolution, proteins were transferred at 30 Volts overnight (18hrs) onto PVDF (Millipore Immobilon-FL, 0.45µM Catalog No. IPFL00010) or nitrocellulose (Bio-Rad Catalog No. 162-0115) with 1X Transfer Buffer (25mM Tris, 192 mM Glycine, 2.5% (v/v) methanol) at room temperature. Transfer efficiency and protein loading was checked via Ponceau S staining and membranes cut into strips based on each protein target’s in-house verified molecular weight. As described previously [107, 108], membranes were blocked with 5% (w/v) milk (Giant, Ellicott City, MD), 0.05% (v/v) Tween-20/1XPBS for 1 hr and subsequently incubated with primary antibody while rocking for 1 hr at room temperature. Following three 10 min washes with PBST (PBS with 0.2% (v/v) Tween-20), blots were incubated with Starbright 520 or 700-conjugated secondary antibodies for 45 min, the membranes were washed thrice for 10 min with PBST, twice for 10 min with PBS and placed on paper towels face-up to dry overnight protected from ambient light. Nitrocellulose-based blots were incubated with horse radish peroxidase-conjugated secondary antibodies for 45 min, the membranes washed as for fluorescent blots, and bands visualized with Clarity Western ECL substrate (BioRad). All blots were imaged using a BioRad ChemiDoc MP imaging system. On occasion, membranes were reprobed with a different primary antibody/secondary antibody combination. In these cases, membranes were re-wet with methanol and subjected to the entire immunoblotting regimen again, but utilizing a secondary antibody conjugated to a different Starbright dye than originally used.

### Reverse transcription-quantitative real time PCR

To determine *PSD1* mRNA levels, yeast were grown overnight at 30°C in 2-4 mL rich lactate. Total RNA was extracted from cell pellets corresponding to 4 OD_600_ using the Quick-RNA Fungal/Bacterial MicroPrep Kit (Zymo Research, Cat. no. R2010) and eluted in 15 µL RNase-free water. To remove residual genomic DNA, RNA samples were treated with 1 µL DNase from the TURBO-DNA free kit (Invitrogen, Cat. No. AM1907), followed by purification with the RNeasy MinElute cleanup Kit (Qiagen, Cat. no. 74204). RNA concentration and purity (A260/A280 ratio ∼2) was assessed with a NanoDrop 8000 spectrophotometer (Thermo). First-strand cDNA was synthesized from 300 ng of total RNA using Superscript VILO Master Mix (Invitrogen, Cat. no. 11754250) according to the manufacturer’s instructions. The quantitative PCR was performed using Applied Biosystems SYBR Green Master Mix in a 20 µl reaction containing 10.5 ng cDNA dilution and 10 µM gene-specific primers. No-template and no-reverse transcriptase (-RT) controls were included in cDNA prep and qPCR. Reactions were run on a Rotor-Gene Q 2plex HRM Platform (Qiagen) and quantification cycle (Cq) values were determined. Relative fold change was calculated using the 2^(-ΔΔCt) method, with *ACT1* as the reference gene. Primer sequences used for qPCR are as follows: *PSD1* forward, 5′-CCAGTAGCACAAGGCGAAGA-3′; *PSD1* reverse, 5′-GACATCAAGGGGTGGGAGTG-3′; and the housekeeping gene *ACT1* forward, 5′-GTATGTGTAAAGCCGGTTTTG-3′; and *ACT1* reverse, 5′-CATGATACCTTGGTGTCTTGG-3′.

### Subcellular fractionation and mitochondrial Isolation

Mitochondrial isolation and subcellular fractionation was carried out as described previously [108]. Precultures of WT and *djp1*Δ strains grown at 30°C for 24-48hrs in rich dextrose or lactate media were used to sterilely inoculate two separate 2L flasks/strain containing 950mL rich dextrose or lactate media with ∼100 OD_600_’s. Final cultures were cultivated at 30°C overnight rotating at 220 rpms, checked for contamination pre-harvesting, and then the cells collected with a 6,000 × *g* centrifugation for 5 min at room temperature. After dispensing the supernatant, the yeast pellets were resuspended with ∼150-200 mL of water, transferred to pre-weighed 250mL bottles, and the cells harvested with a 2000 × *g* centrifugation for 5 min at room temperature. After decanting the supernatant and weighing the yeast pellets, cells were resuspended in 50mL of newly made 0.1M Tris-SO_4,_ pH 9.4 (1M Tris with pH adjusted with sulfuric acid) containing 15mM dithiothreitol, and incubated shaking at 220rpm for 20 min at 30°C. Yeast were next collected with a 2000 × *g* centrifugation for 5 min at room temperature, washed with 40mL 1.2M sorbitol, 20mM KPi, pH 7.4 buffer, and re-collected as before. Upon discarding the supernatant, yeast were converted to spheroplasts by resuspending in 1.2M sorbitol, 20mM KPi, pH 7.4 (2mL per gram of yeast) containing Zymolyase 20T (Nacalai Tesque, INC.; 3mg per gram of yeast) and incubating for ∼1hour at 30°C shaking at 220 rpm. Digestion of the cell wall was assessed by checking for yeast lysis upon dilution in 10µL water. Spheroplasts were collected with a 3500 × *g* centrifugation for 5 min at 4°C and the resulting supernatant discarded; all subsequent steps were performed at 4°C and/or on ice. Following two washes with 1.2M sorbitol, 20mM KPi, pH 7.4 buffer chilled at 4°C, with the spheroplasts collected by a 3500 × *g* spin for 5 min at 4°C after each wash, the pellet was resuspended in 50mL of 0.6M sorbitol, 20mM KOH-MES, pH 6.0 (BB6.0 buffer) containing 1mM phenylmethylsulfonyl fluoride (PMSF), poured into a tight-fitting (type A) glass dounce homogenizer kept on ice, and then homogenized using 15 strokes. 200µL of the resulting homogenate was set aside to quantitate and analyze as the starting material (SM). Following a 4°C, 1700 × *g* 5 min spin, the supernatant was set aside and the residual pellet containing unbroken cells re-homogenized in BB6.0 + 1mM PMSF by 15 strokes in the same glass dounce to extract more membrane-bound organelles. The resulting homogenate was centrifuged as before and the supernatants from both homogenizations combined and centrifuged for 10 min at 13,500 × *g* and 4°C. The supernatant (S13) was either side aside for additional fractionation steps or thrown out whereas the mitochondrial-enriched pellet was washed with 35mL of BB6.0 buffer bereft of PMSF. Mitochondria were resuspended by two up-and-down strokes with a pre-chilled Teflon dounce and after a 1700 × *g* 5 min spin at 4°C, the mitochondria-containing supernatant was transferred to a fresh 50mL tube while the pellets were discarded. Mitochondria were again collected with a 13,500 × *g* centrifugation for 10min at 4°C after which the supernatant was removed by aspiration and the mitochondrial pellet homogenized by two up-and-down strokes in 30mL of BB7.4 buffer with the same pre-chilled Teflon dounce as before. After a final 13,500 × *g* spin for 10min at 4°C in a fresh 50 mL tube, the supernatant was aspirated away from the mitochondrial pellet (P13), which was resuspended in leftover BB7.4 and homogenized by one stroke in a small pre-chilled Teflon dounce prior to determining the protein concentration of the mitochondrial slurry using the Pierce BCA Protein Assay Kit (ThermoFisher Scientific, Catalog No. 23225). Mitochondria were snap-frozen in liquid nitrogen in convenient aliquots, typically 1mg/tube, and stored at -80°C.

To harvest additional subcellular fractions, 35mL of the S13 supernatant was transferred to a 50mL tube and centrifuged at 21,500 × *g* for 15 min at 4°C. Post-spin, the supernatant was transferred to a fresh 50mL tube and spun at 40,000 × *g* for 30min at 4°C. The 40,000 *x g* supernatant (S40) was transferred to a 50mL falcon tube and the 40,000 × *g* pellet (P40) was resuspended in residual buffer and transferred to a 1.5mL microcentrifuge tube on ice. The protein concentration of each fraction (SM, P13, P40, and S40) was determined with the BCA assay and aliquots of each snap frozen in liquid nitrogen and stored at -80°C.

#### Protein structure rendering

Representative protein structure images were generated from available AlphaFold models using the biomolecular visualization software ChimeraX (version 1.12).

### TMT Proteomics

#### Sample preparation

Proteins were isolated from purified mitochondria (P13 fractions shown in Figure 6) using SP3 paramagnetic beads (GE Healthcare) [109]. Briefly, 10 µg of mitochondria was solubilized in 10mM TEAB and disulfide bonds reduced with dithiothreitol (5 mM final concentration) for 1 hour at 60°C. Samples were cooled to room temperature and pH adjusted to ∼8.0, followed by alkylation with iodoacetamide (10 mM final concentration) in the dark at room temperature for 15 minutes. Next, SP3 beads were added to the samples at a 10:1 bead:protein ratio and 100% acetonitrile was added to achieve a final acetonitrile concentration of 50% (v/v). Samples were incubated at room temperature with shaking for 5 minutes. Following protein binding, beads were washed with 180 µL 80% ethanol three times. Proteins were digested on-bead with trypsin (Pierce) at 37°C overnight at a 50:1 protein:enzyme ratio and peptides were removed from beads and dried by vacuum centrifugation.

#### Isobaric tandem mass tag (TMT) labeling and peptide fractionation

Peptides were labeled with TMTpro 16-plex reagents (Thermo Fisher) according to the manufacturer’s instructions and the reaction was quenched with hydroxylamine. All 6 TMT-labeled peptide samples were combined and dried by vacuum centrifugation. The pooled TMT-labeled sample was re-constituted in 100 µL 200mM TEAB buffer and desalted with Pierce Detergent removal columns (Fisher Scientific PN 87777) to remove excess TMT label. Peptides in the eluate (∼300 µL) were diluted to 2 mL with 10 mM TEAB and fractionated on a XBridge C18 Column (5 µm, 2.1 x 100 mm column (Waters) and XBridge C18 Guard Column (5 µm, 2.1 x 10 mm, Waters) using a 0 to 90% acetonitrile in 10 mM TEAB gradient over 85 min at 250 µL/min on an Agilent 1200 series capillary HPLC with a micro-fraction collector. Eighty-four 250 ul fractions were collected and concatenated into 24 fractions according to Wang et al 2011 [110] and dried. *Liquid chromatography separation and tandem mass spectrometry (LCMS/MS)*.

Dried fractions were reconstituted in 2% acetonitrile/0.1% formic acid and analyzed by nanoflow liquid chromatography-tandem mass spectrometry (nLC-MS/MS) using a Vanquish UHPLC interfaced with an Orbitrap Exploris 480 mass spectrometer (Thermo Fisher Scientific). Peptide separation was performed with a 120-minute linear water/acetonitrile gradient over polyimide-coated, fused-silica, 25 cm × 360 μm o.d./75 μm i.d. self-packed PicoFrit columns (New Objective) with built-in emitters (75 µm emitter i.d.). Stationary phase in analytical columns consisted of ReproSil-Pur 120 C18-AQ, 2.4 μm particle size, 120 Å pore (Dr. Maisch High Performance LC GmbH). Trap columns consisted of ∼1 cm × 360 μm o.d./75 μm i.d. polyimide-coated, fused-silica tubing (New Objective), packed with 5 μm particle size, 120 Å pore, C18 stationary phase (ReproSil-Pur), with a Kasil frit. Electrospray ionization was accomplished with 2 kV positive spray voltage and an ion transfer tube temperature of 250°C.

MS1 precursor ion scans were acquired from 375-1500 m/z (rich dextrose mitochondria) or 350-1800 m/z (rich lactate mitochondria) at 120,000 resolution at 200 m/z with a normalized AGC of 300%, an RF lens setting of 50%, and maximum injection time set to Auto. Precursor ions were isolated with a 0.7 m/z isolation window and the following data-dependent filters: monoisotopic precursor selection (MIPS) set to peptide mode, intensity threshold 5×104, charge states 2-6, dynamic exclusion duration of 45 s (rich dextrose mitochondria) or 15 s (rich lactate mitochondria; max n=1 scan, 10 ppm tolerance), and Advanced Peak Determination activated. Precursors were fragmented by HCD with a 34% (rich dextrose mitochondria) or 32% (rich lactate mitochondria) normalized collision energy and MS/MS spectra were acquired in TurboTMT mode at 30,000 resolution (rich dextrose mitochondria) or at 45,000 resolution (rich lactate mitochondria) with AGC set to Standard and maximum injection time set to Auto. The duty cycle was restricted to a maximum of 3 seconds (rich dextrose mitochondria) or 15 MS/MS scans (rich lactate mitochondria) between MS survey scans.

#### Data analysis

The raw data from rich lactate mitochondria was searched with Mascot (version 2.8) in Proteome Discoverer (version 2.5) against an *S. cerevisiae* RefSeq FASTA database (5,555 entries). Search criteria were tryptic cleavage (maximum 1 missed), 3 ppm precursor ion tolerance, 0.01 Da fragment ion mass tolerance, Cys carbamidomethylation and TMTpro on peptide N-termini as fixed modifications, and TMTpro on Lys, deamidation on Asn/Gln, and Met oxidation as variable modifications. Peptide identifications were validated by Percolator at 5% false discovery rate (FDR) using an auto-concatenated target-decoy approach. Peptide spectral matches (PSMs) were filtered for ≤30% precursor ion isolation interference and only proteotypically unique peptides were used for relative quantitation. TMT reporter ion abundances were normalized to the total summed abundance per channel and quantification was based on reporter ion signal-to-noise values. Group differences were tested by ANOVA based on protein abundance with no imputation of missing values. The raw data from rich dextrose mitochondria was searched with Chimerys (Inferys model 3.0) in Proteome Discoverer (version 3.2) against an *S. cerevisiae* UniProt FASTA database (5,993 entries) and a database containing common contaminants (438 entries). Search criteria were tryptic cleavage (maximum 1 missed), 20 ppm fragment ion mass tolerance, Cys carbamidomethylation, TMTpro on peptide N-termini, and TMTpro on Lys as fixed modifications and Met oxidation as a variable modification. Peptide identifications were validated by Chimerys at 1% false discovery rate (FDR) using an auto-concatenated target-decoy approach. Peptide spectral matches (PSMs) were filtered for ≤ 30% precursor ion isolation interference and ≥0.8 Chimerys coefficient. TMT reporter ion abundances were normalized to the total summed abundance per channel and quantification was based on reporter ion signal-to-noise values. Group differences were tested by ANOVA based on protein abundance with no imputation of missing values.

#### Cellular Compartment Assignment

Cellular compartment for all proteins was determined using the Bioconductor R package org.Sc.sgd.db (R version 4.6.0). Cellular compartment designation for all proteins with significant (p-value ≤ 0.05) log2FC values was cross-referenced with the cellular compartment information available on the Saccharomyces Genome Database (SGD). The Locus Overview summary, Gene Ontology summary, and Gene Ontology cellular compartment sections were checked for the words “mitochondrion”, “mitochondria”, “mitochondrial”, “peroxisome”, and “peroxisomal”. Any mention of these cellular compartments for a given gene in SGD where the cellular compartment designation had not already been assigned by the R package was manually designated as such.

#### Volcano Plots and Heatmaps

Volcano plots were generated using the ggplot2 package in R (R version 4.6.0). Volcano plot points were colored according to their cellular compartment assignment for all proteins with significant (p-value ≤ 0.05) log2FC values. Any proteins with non-significant (p-value > 0.05) log2FC values were colored gray, regardless of cellular compartment localization. Proteins with mitochondrial localization were colored red. Proteins with peroxisomal localization were colored light blue. Proteins with both mitochondrial and peroxisomal localization were colored purple. Any proteins with significant (p-value ≤ 0.05) log2FC values, but with neither mitochondrial nor peroxisomal localization, were colored black. Heatmaps were generated using the pheatmap package in R (R version 4.6.0). Proteins were included in heatmaps if the protein met the following inclusion criteria for at least one of the sample groups: p-value ≤0.05 and log2FC < -0.38 or > 0.38. Any protein that met the log2FC inclusion criteria for only one sample group was colored gray for the sample group in which the protein did not meet inclusion criteria and any protein that was not included in the dataset for one of the sample groups was colored black. All proteins that had log2FC with p-value ≤0.05 are labeled with an asterisk (*).

### Lipid Steady States

After 2-3 days growing at 30°C with 220 rpm shaking in 2 mL of rich lactate medium, starter cultures were used to inoculate 2mL SD for final OD_600_ = 0.09 (based on delayed growth, a final OD_600_ = 0.36 was used for *psd1*Δ*djp1*Δ strains), 2 mL SD spiked with 2mM ethanolamine for final OD_600_ = 0.09 (final OD_600_ = 0.36 and 0.18 for *psd1*Δ*djp1*Δ and for *psd2psd1*Δ*djp1*Δ strains, respectively), 2 mL SDLac for final OD_600_ = 0.18 (final OD_600_ = 0.5 for *psd1*Δ and *psd1*Δ*djp1*Δ strains, respectively), and 2 mL SDLac with 2mM ethanolamine for final OD_600_ = 0.18 (final OD_600_ = 0.36 for *psd1*Δ*, psd1*Δ*djp1*Δ, *psd2psd1*Δ, and *psd2psd1*Δ*djp1*Δ strains, respectively). 1µCi ^14^C-acetate (Perkin Elmer, NEC084H001MC) was flame-sterilely added to each sample to initiate lipid labeling and all of the cultures were grown in a shaking (240 rpm) water bath at 30°C for 24 hrs. The next day, yeast were collected with a 3000 rpm spin in a clinical centrifuge for 5 min at room temp. The radioactive supernatants were then aspirated, the pellets washed with 2mL sterile water, and the yeast collected as before. Post-aspiration, yeast pellets were resuspended with 0.3mL MTE buffer (0.65M Mannitol, 20mM Tris, 1mM EDTA) spiked with protease inhibitors (1mM PMSF, 10µM leupeptin, and 2µM pepstatin A), transferred to 1.5mL microcentrifuge tubes containing ∼ 0.1mL glass beads, parafilm-sealed, and the contained yeast mechanically destroyed by vortexing for 35 min on high in a cold room. After a brief 2-min, 4°C spin at 376 × *g*, the supernatants were transferred to clean 1.5mL microcentrifuge tubes and crude mitochondria collected with a 13,000 × *g* centrifugation for 5 minutes at 4°C. Once the supernatants were removed by aspiration, the mitochondrial-enriched pellets were resuspended by repeat pipetting in 50µL BB7.4 (0.6M Sorbitol, 20mM HEPES-KOH, pH 7.4). ^14^C- acetate incorporation was then measured in 1µL of mitochondrial homogenate by liquid scintillation counting. Prior to lipid extraction, ∼0.5mg cold mitochondria was added to each sample to improve eventual lipid migration on thin layer chromatography (TLC) plates. Phospholipids were extracted from equal amounts of labeled mitochondria (based on ^14^C-acetate incorporation) in 5mL borosilicate tubes using 1.5mL 2:1 Chloroform:Methanol and vortexing on medium-high for 30 minutes at room temperature. Phase separation was initiated by adding 0.3mL of Normal Saline (0.9% (w/v) NaCl) and the samples vortexed for an additional 1 min. Samples were centrifuged (Sorvall Legend X1R) at 1000 rpm for 5 min at room temperature and the upper aqueous phase aspirated. The remaining organic phase was washed with 0.25mL 1:1 Methanol:H_2_O, vortexed for 30 sec, and phases separated as before. The lower organic phase was carefully transferred using a short Pasteur pipette controlled by a Pipette Pump^TM^ (SP Bel-Art) to a new 5mL borosilicate tube, dried down under a stream of nitrogen, and either analyzed directly by TLC or stored at -20°C until ready to resolve. TLC plates (Machery-Nagel SILGUR 25) were washed with chloroform, air-dried, pre-treated with 1.8% (w/v) boric acid in 100% ETOH, and activated at 95°C for at least 30 min. Dried lipid samples were resuspended in 40µL chloroform and 13µL of each loaded onto TLC plates using a CAMAG Linomat 5. Plates were resolved in equilibrated TLC tanks containing 30:35:7:35 (v/v) chloroform:ethanol:water:triethylamine, air dried, exposed to phosphorimaging K screens, and signals detected using a Molecular Dynamics Storm 820 Phosphor Imager.

### Recombinant Proteins and Antibodies

The custom mouse monoclonal antibody against yeast Crd1 (8G12F4) was produced by Genscript and custom rabbit antisera against yeast Ggc1 (3673.3), yeast Gep4 (7296.F), yeast Msp1 (20541.1), and yeast Get3 (20543.2) were generated by Pacific Immunology (Ramona, CA) using affinity-purified His_6_Crd1, His_6_Ggc1, His_6_Gep4, His_6_Msp1, and His_6_Get3 as antigens. The specificity of these custom-made antibodies are documented in Fig S1. In brief, the predicted mature open reading frame for Crd1 (Pro-57) and the entire open reading frames of Ggc1, Gep4, Msp1, and Get3 were cloned downstream of the His_6_ tag encoded in the pET28a plasmid (Novagene) and transformed in BL21(RIL) *Escherichia coli*. Each protein was induced with 0.5 mM IPTG in 1L 2X YT cultures containing 20µg/mL kanamycin for 4 h at 25°C and the bacterial pellet collected at 3020 × *g* for 10 min, washed with 0.9% (w/v) NaCl, and stored at −20°C. For His_6_Ggc1, the pellet was thawed on ice and resuspended in 30mL BPER II reagent (Thermofisher 78260) with a transfer pipet interspersed with vortexing. Once homogenized, the mixture was shaken by hand at room temperature for 10min and then centrifuged at 27,000 × *g* for 15min at room temperature. The resulting supernatant was set aside whereas the pellet was again resuspended with 30mL BPER II, and once accomplished, treated with lysozyme (0.2 mg/mL final) for 5min at room temperature before 100mL 1:20 diluted BPER II (diluted with ddH_2_0) was added and the inclusion bodies collected at 35,000 × *g* for 15min at 4°C. The supernatant was decanted and the pellet resuspended with 100mL 1:20 diluted BPER II by vortexing and again spun at 35,000 × *g* for 15min at 4°C. This washing step was repeated 2 additional times. The final pellet was vortexed on high with 5mL Inclusion Body solubilization buffer (1.67%(w/v) Sarkosyl, 0.1mM EDTA, 10 mM Tris-Cl pH 7.4, 0.05% (w/v) PEG3350) freshly spiked with 10 mM DTT and once homogenized, incubated on ice for 20 min and then diluted with 10mL 10 mM Tris-Cl pH 7.4. His_6_Ggc1 (4mL) was diluted with 34mL10 mM Tris-Cl pH 7.4, 0.01% (w/v) PEG3350, 0.1% (v/v) glycerol), purified by incubating with 1.5 mL Ni-NTA in a 50mL falcon tube with rotation for 2hr at 4°C, and the flow through collected after being loaded into a chromatography column. The Ni-NTA resin was sequentially washed with 10mL Buffer C (100 mM NaH_2_PO_4_•H_2_O, 10 mM Tris-Cl, 8 M Urea, pH 6.3) and 10mL Buffer D (100 mM NaH_2_PO_4_•H_2_O, 10 mM Tris-Cl, 8 M Urea, pH 5.9), and the bound His_6_Ggc1 thrice-eluted with 0.5mL reducing sample buffer lacking bromophenol blue (20 mM Tris-Cl pH 7.4, 2% (w/v) sodium dodecyl sulfate, 10% (v/v) glycerol, 25mM DTT). The recovered His_6_Ggc1 was dialyzed against 20mM HEPES-KOH pH 7.4, 150mM NaCl, 1mM MgCl_2_, 0.1% (w/v) sarkosyl) and quantified against a BSA standard curve.

For His_6_Crd1, His_6_Gep4, His_6_Msp1, and His_6_Get3, the frozen bacterial pellets were thawed and then resuspended in 40mL lysis buffer (50mM NaH_2_PO_4_, 300mM NaCl, 10mM Imidazole, 0.1mM EDTA, pH 8.0) containing 1mg/mL lysozyme and either 1% (v/v) Tween-20 (His_6_Crd1 and His_6_Gep4) or 0.05% (v/v) Tween-20 (His_6_Get3 and His_6_Msp1) and incubated with rocking for 30 min at 4°C. The resulting bacterial suspensions were emulsified using an Avestin Homogenizer and the resulting lysate centrifuged at 10,000 × *g* at 4°C for 20 min. His_6_Get3 was enriched in the resulting supernatant (cell extract) whereas His_6_Crd1, His_6_Gep4, and His_6_Msp1 largely fractionated in the resulting pellet. The His_6_Crd1-, His_6_Gep4-, and His_6_Msp1-enriched pellets were dispersed with 5mL Inclusion Body solubilization buffer (1.67%(w/v) Sarkosyl, 0.1mM EDTA, 10 mM Tris-Cl pH 7.4, 0.05% (w/v) PEG3350) freshly spiked with 10 mM DTT by vortexing on high and then incubating on ice for 20 min before being diluted with 10mL 10mM Tris-Cl pH 7.4. His_6_Get3 in the cell extract (∼40mL) or inclusion body-extracted His_6_Crd1, His_6_Gep4, and His_6_Msp1 (∼15mL) were purified by incubating with 1.5 mL Ni-NTA in a 50mL (His_6_Get3) or 15mL (His_6_Crd1, His_6_Gep4, and His_6_Msp1) falcon tube. After rotating for 2hr at 4°C, the Ni-NTA mixtures were loaded onto chromatography columns and the non-binding flow throughs collected. The His_6_Crd1 and His_6_Gep4 columns were washed first with 14mL of 0.1% (w/v) Sarkosyl, 50mM NaH_2_PO_4_, 300mM NaCl, 20 mM Imidazole, 10% (v/v) glycerol, pH 8.0 and then with 14mL of 0.1% (w/v) Sarkosyl, 50mM NaH_2_PO_4_, 600mM NaCl, 20 mM Imidazole, 10% (v/v) glycerol, pH 7.0. The His_6_Get3 and His_6_Msp1 columns were subjected to four 10mL washes with 1) 0.1% (w/v) Sarkosyl, 50mM NaH_2_PO_4_, 300mM NaCl, 20 mM Imidazole, 10% (v/v) glycerol, 20mM β-ME, pH 8.0; 2) 0.15% (w/v) Sarkosyl, 50mM NaH_2_PO_4_, 600mM NaCl, 30mM Imidazole, 20% (v/v) glycerol, 20mM β-ME, pH 8.0; 3) 0.2% (w/v) Sarkosyl, 50mM NaH_2_PO_4_, 600mM NaCl, 40mM Imidazole, 30% (v/v) glycerol, 20mM β-ME, pH 8.0; and 4) 0.1% (w/v) Sarkosyl, 50mM NaH_2_PO_4_, 300mM NaCl, 20 mM Imidazole, 10% (v/v) glycerol, 20mM β-ME, pH 8.0. Bound His_6_Crd1, His_6_Gep4, His_6_Msp1, and His_6_Get3 was then recovered with elution buffer (250mM imidazole, 0.1% (w/v) Sarkosyl, 50mM NaH_2_PO_4_, 300mM NaCl, and 10% (v/v) glycerol, pH 8.0; 6 sequential 0.5mL elutions). Protein-containing fractions were identified using the Bradford Assay (Bio-Rad) and additional elutions performed as needed until the resulting eluates were devoid of detectable protein as compared to the buffer blank. Protein-containing eluates were combined, PBS dialyzed, and quantified using a BSA standard curve prior to antibody generation.

Other in-house antibodies generated in either in our laboratory or the laboratory of C. Koehler (UCLA) and used in this study include rabbit anti-yeast Psd1 (Psd1β-specific; 4077.5 and 4078.5; 1:1000; [111]), rabbit anti-yeast Pic1 (3676.3; 1:10,000; [112]), rabbit anti-yeast Mic60 (20450.F; 1:5000; [25]), rabbit anti-Msp1 (20541.1; 1:1000; this study), rabbit anti-Gep4 (7926.F; 1:2000, this study), mouse anti-Crd1 (8G12F4; 1:1000; this study), rabbit anti-Cld1 (5480.T; 1:1000; [105]), rabbit anti-yeast Taz1 (4248.F; 1:1000; [108]), rabbit anti-yeast Por1 (425; 1:10,000; [113]), rabbit anti-yeast Tom70 (7306.F; 1:10,000; [114]), rabbit anti-yeast Hsp70 (SH1-T; 1:10,000; [108]), rabbit anti-Ggc1 (3673.3; 1:2000; this study), rabbit anti-yeast Tim50 (3543.T; 1:1000; [115]), and rabbit anti-yeast Yme1 (αYme1; 1:1000; [116]). Additional antibodies employed were rabbit anti-yeast Djp1 (αDjp1; 1:2000; [41]), rabbit anti-yeast Mic60 (αFcj1; 1:1000; [117]), mouse anti-FLAG (clone M2; 1:5000; Sigma F3165;), mouse anti-His (1B7G5; 1:3000-5000; ProteinTech 66005-1), mouse anti-Dpm1 (5C5A7; 1:1000; Abcam 113686), mouse anti-yeast Aac2 (6H8; 1:1000; [118]), mouse anti-yeast Sec62 (Blue top; 1:1000; David Meyer), mouse anti-Myc (Myc.A7; 1:2000; Thermofisher MA1-21316), and Starbright 520/700-conjugated (BioRad; all immunoblots except for Figure 2D) and horseradish peroxidase-conjugated (Thermofisher; Figure 2D only) secondary antibodies.

### Statistical Analyses

Tif images of the immunoblots and TLC plates were quantitated by Quantity One or ImageLab (BioRad Laboratories) and statistical comparisons (ns, *P >* 0.05; 1 symbol *P ≤* 0.05; 2 symbols *P ≤* 0.01; 3 symbols *P ≤* 0.001; 4 symbols *P ≤* 0.0001) performed using Prism 11 (GraphPad). All graphs depict the mean ± SD. The sample size and statistical test performed are indicated in the associated figure legends.

#### Miscellaneous

All presented immunoblots and TLC images are representative of at least three independent experiments performed on three separate days.

## Acknowledgements

We would like to thank Drs. Mike Renne (Saarland University, Germany) for SD yeast media formulations and Carla Koehler (UCLA, USA) and Johannes Herrmann (University of Kaiserslautern) for antibodies. The TMT proteomics was done by the Mass Spectrometry and Proteomics Facility in the former Biological Chemistry Department (rich lactate mitochondria) and the Center for Proteomics Discovery in the Institute for Cell Engineering (rich dextrose mitochondria), both at Johns Hopkins University.

This manuscript was supported by the National Institutes of Health (NIH; NIH grants R01GM151746, R01GM111548, and R01GM111548-08S2 to SMC; R01GM111548-03S1 to PNS; and R01GM111548-07S1 to AMR) and the National Science Foundation Graduate Research Fellowship Program (DGE1746891 to PNS). The content is solely the responsibility of the authors and does not necessarily represent the official views of the National Institutes of Health. It is subject to the NIH Public Access Policy. Through acceptance of this federal funding, NIH has been given a right to make this manuscript publicly available in PubMed Central upon the Official Date of Publication, as defined by NIH.

---

**Fig S1.**
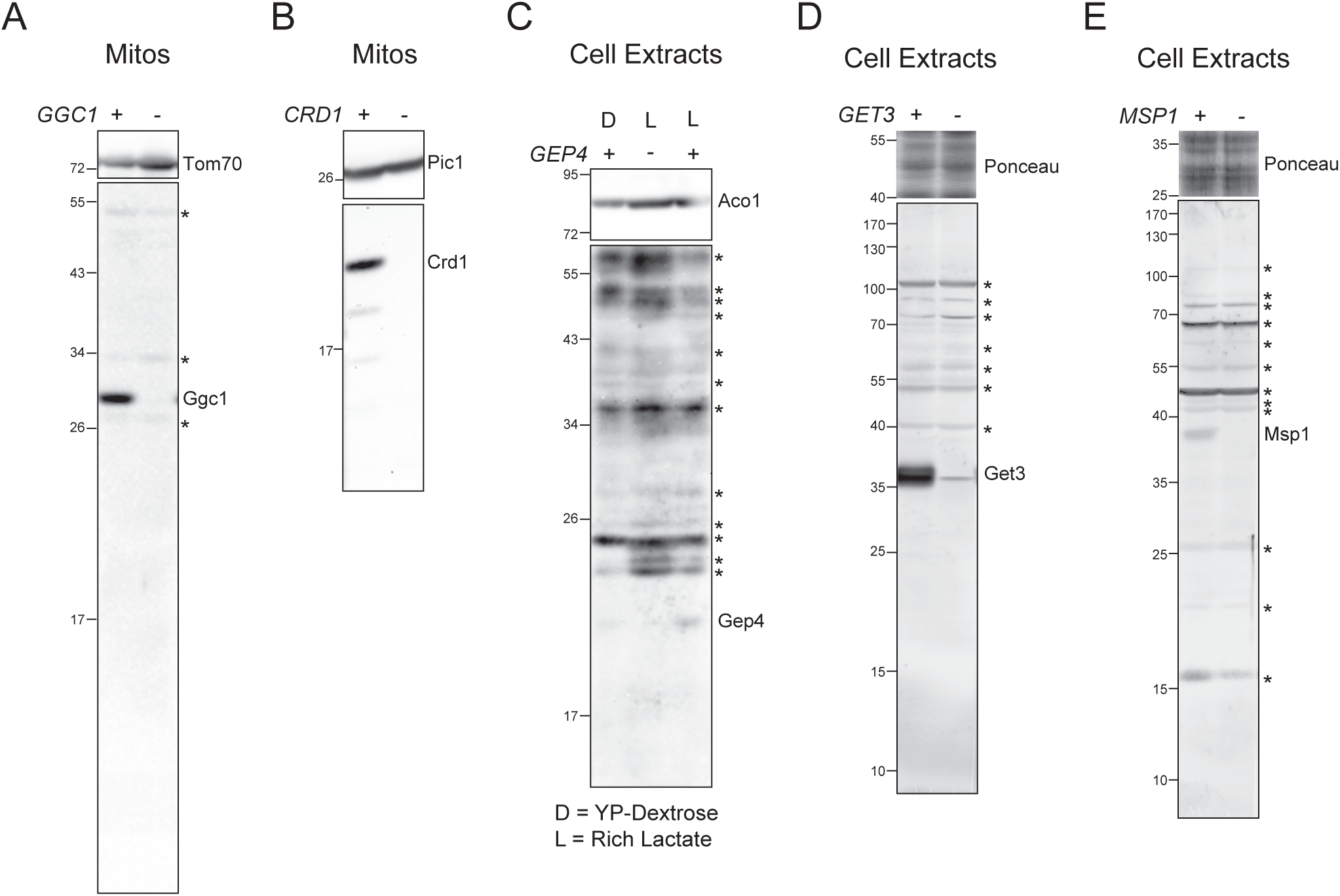
Custom antibody specificity. **(A)** 25µg mitochondria from indicated yeast strains were immunoblotted for Ggc1; Tom70 served as loading control. **(B)** 40µg mitochondria from indicated yeast strains were immunoblotted for Crd1; Pic1 served as loading control. **(C)** Cell extracts derived from the designated strains grown at 30°C in the indicated rich media were immunoblotted for Gep4; Aco1 served as loading control. **(D)** Cell extracts derived from the indicated strains grown at 30°C in rich lactate media were immunoblotted for Get3; total protein stain served as loading control. **(E)** Cell extracts derived from the indicated strains grown at 30°C in rich lactate media were immunoblotted for Msp1; total protein stain served as loading control.

